# Pronounced genetic structure associated with differences in a reproductive trait and climatic barriers in Canadian populations of the western toad (*Anaxyrus boreas*)

**DOI:** 10.1101/2025.06.17.660222

**Authors:** Jayna C. Bergman, Juan Enciso-Romero, Gregory B. Pauly, Roseanna Gamlen-Greene, Melissa Todd, Julie A. Lee-Yaw

## Abstract

Identifying genetic groups within species is critical for inventorying biodiversity, understanding species’ distributions, and delineating conservation units. The western toad (*Anaxyrus boreas*) is one of the most widespread amphibians in western Canada and one of just two amphibian species to maintain an extensive range on both sides of the Canadian Rocky Mountains. Curiously, toads on the eastern side of the mountains have a vocal sac and produce a distinct advertisement call whereas toads on the western side of the mountains lack these traits. However, the extent to which these “Calling” and “Non-Calling” populations are genetically and ecologically distinct remains unclear. We used reduced representation sequencing (ddRAD-seq) to genotype individuals from across the Canadian portion of the species’ range. Combining population genomic analysis with the original phenotypic data used to delineate the Calling and Non-Calling populations and with predictions from ecological niche models, we show that Calling and Non-Calling populations of western toads represent distinct genetic groups separated by both differences in a major reproductive trait and by climate barriers. We additionally find evidence for a third, undescribed group in southwestern Alberta and southeastern British Columbia. Our results suggest Calling and Non-Calling populations should be managed as distinct groups and highlight the potential for there to be cryptic and strong genetic structure among northern populations of widespread species.

## Introduction

Characterizing intraspecific genetic structure is fundamental to understanding species’ geographic distributions. In particular, genetic structure provides insight into the history of species’ ranges and the factors influencing colonization (Hewit, 2004; Cairns et al., 2021), as well as the extent to which populations are currently connected. Genetic breaks in the range inform our understanding of geographic and ecological barriers that influence dispersal (Waters et al., 2020; Suarez et al., 2022) and thus our understanding of the ability of species to track suitable conditions as the climate changes. Finally, assessments of genetic structure within species’ ranges are important for inventorying biodiversity, allowing for the identification of cryptic taxa (Bickford et al., 2007; Fiser et al., 2017) and the delineation of conservation units for wildlife management (Funk et al., 2012; Coates et al., 2018; Forester et al., 2022).

Genetic substructure is expected to be particularly pronounced in the Northern Hemisphere, where species experienced repeated range expansions and contractions during the Pleistocene glaciations (Hewitt, 2004). Indeed, numerous studies have used molecular markers and methods to assess substructure within species’ ranges in areas impacted by the Pleistocene glaciations, revealing distinct genetic groups within what are otherwise widespread species (see review by: Soltis et al., 1997; Hewitt, 2000; Brunsfelt et al., 2001; Sommer and Zachos, 2009; Shafer et al., 2010; Lyman and Edwards, 2022; Fu and Wen, 2023; Garcia-Rodriguez et al., 2024). However, gaps remain in our understanding of structure in some regions. For example, for some taxonomic groups in North America, genetic studies are heavily biased towards populations in the United States, limiting our understanding of genetic structure in the previously glaciated, northern portion of many species’ ranges (Lesbarrères et al., 2014; Bergman et al. unpublished). In other cases, what is known about genetic structure across Canada comes from earlier studies using mitochondrial markers or a limited number of nuclear markers (e.g. Graham and Burg, 2012; Benefer et al., 2013; Nobarinezhad et al., 2020; but see Lehnert et al., 2023; Miller et al., 2024; Stacy et al., 2025). It is now widely recognized that issues with incomplete lineage sorting, resolution, and/or introgression may limit the conclusions that can be drawn from studies using sparse sampling of the genome to assess genetic structure (e.g. Ballard and Whitlock, 2004; Funk et al., 2019). Thus, for many northern species, our understanding of intraspecific genetic diversity remains incomplete.

Next generation sequencing has provided an opportunity to revisit questions about genetic structure within species’ ranges using information from across the genome and has clarified the number and distribution of genetic groups within the ranges of several species (e.g. Barbosa et al., 2018; Dinca et al., 2019; Cairns et al., 2021; Gallego-García et al., 2021). However, care is needed when using genomic data to delineate meaningful groups. For instance, when there are gaps in the sampling of the geographic range, what is continuous genetic variation associated with isolation-by-distance (IBD) may be misinterpreted as distinct genetic substructure (Chambers and Hillis, 2020; Turbek et al., 2023). Ensuring samples are representative of the transition across suspected contact zones is thus important for identifying true genetic breaks within species’ ranges (Chambers et al., 2022; Turbek et al., 2023). At the same time, the high resolution of genomic data is likely to reveal fine-scale structure, which may lead to issues with the over splitting of groups (Sukumaran and Knowles, 2017; Coates et al., 2018; Chambers et al., 2025). Thus, it is ideal to pair genomic assessments of structure with phenotypic (e.g. Gordon et al., 2020; Waples and Lindley, 2018) and/or ecological (e.g. Campbell et al., 2022) data that can provide insight into the biological significance of any observed groups.

Amphibians are expected to exhibit strong genetic structure within their ranges because they have low dispersal ability, are often philopatric, and show evidence of limited gene flow over small spatial scales (Beebee, 2005; Zeisset and Beebee, 2008). Indeed, amphibians in North America have been shown to consist of distinct genetic groups, with continentally-distributed species often consisting of eastern and western clades (e.g. Lee-Yaw et al., 2008; Everson et al., 2021). Within the eastern United States, intraspecific genetic structure is often aligned with major geographic features such as the Atlantic Coast of the Florida peninsula, Apalachicola and Mississippi Rivers, and the Appalachian Mountains (Soltis et al., 2006; Pauly, et al., 2007; Lyman and Edwards, 2022). In the west, sharp genetic breaks also tend to line up with major geographic features, including the Cascade/Coast mountains range, the southern Rocky Mountains, and the Columbia River Basin (Zeisset and Beebee, 2008; Shafer et al., 2010). However, most phylogeographic studies of North American amphibians are based on populations in the United States (Lesbarrères et al., 2014; but see Lee-Yaw et al., 2008; Wilson et al., 2008; Lee-Yaw and Irwin, 2012; Cairns et al., 2021), limiting our understanding of the features that shape genetic structure across much of North America, as well as efforts to protect genetic diversity in northern areas.

Western toads (*Anaxyrus boreas*; also known as *Bufo boreas*) are one of the most widely distributed amphibians in western North America and are one of just two amphibian species in Canada to maintain an extensive range on either side of the Canadian Rocky Mountains (CRMs), a barrier that shapes genetic diversity in other taxonomic groups (Shafer et al., 2010; Jensen et al., 2024). Furthermore, western toads on either side of the CRMs exhibit differences in key reproductive traits: individuals east of the mountains in Alberta have a vocal sac and pronounced advertisement call, whereas those to the west in British Columbia (BC) lack both traits (Fig. 1; Pauly, 2008). To the best of our knowledge, this is the only documented case in anurans where there is a polymorphism in terms of the presence of and ability to produce male advertisement calls within the range of the same species (see Elias-Costa & Faivovich 2025 for more general discussion of the gain and loss of calls across Anurans). Based on these differences, “Calling” and “Non-Calling” populations are recognized as distinct conservation units in Canada (i.e. Designatable Units) by the Committee on the Status of Endangered Wildlife in Canada (COSEWIC 2012). Yet, despite attention to genetic structure elsewhere in the species’ range (Goebel et al., 2009; Switzer et al., 2009; Addis et al., 2015; Gordon, 2017; Gordon et al., 2020; US Fish and Wildlife, 2017; Gamlen-Greene, 2022; Trumbo et al., 2023; Fig. 1A), the extent to which Calling and Non-Calling populations of western toads are genetically distinct remains unclear. Given conservation concern for western toads across much of their range (e.g. Colorado Parks and Wildlife, New Mexico Department of Game and Fish, Wyoming Department of Game and Fish), including in Canada (COSEWIC, 2012), clarifying genetic structure in this part of the species’ range not only adds to our understanding of genetic diversity in the north but has the potential to guide management plans for the species.

**Figure 1.**
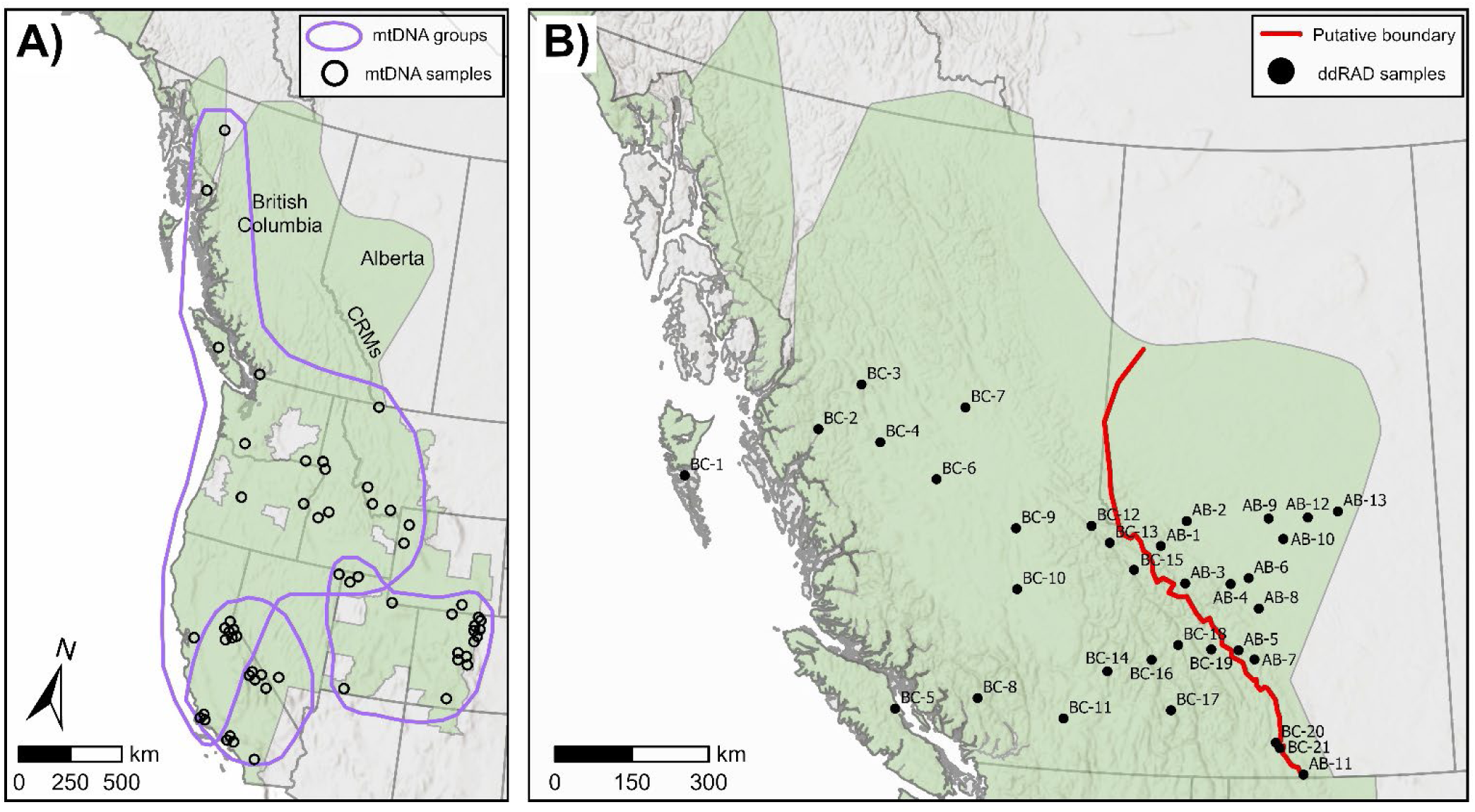
(A) The geographic range of western toads (*Anaxyrus boreas*) in western North America (green shading). Open points represent approximate locations of the sampling sites used to identify mitochondrial groups (purple polygons) from previous work (recreated from Goebel et al., 2009). Genetic structure across the Canadian range, including across the Canadian Rocky Mountains (CRMs) is unknown. (B) Locations of the genetic sampling sites for the present study. The putative boundary between the Calling and Non-Calling populations is shown as a red line (redrawn from: Environment and Climate Change Canada, 2020).

In this study, we evaluate genetic structure across the Canadian range of western toads. We specifically ask whether genomic data support the recognition of the Calling and Non-Calling populations as distinct groups and whether observed genetic boundaries line up with previously described variation in call morphology (Pauly 2008). We additionally use ecological niche models to explore niche overlap between Calling and Non-Calling toads and the potential for gene flow between groups. We discuss the implications of our results for the conservation status of the Calling and Non-Calling populations in Canada and for our broader understanding of the biogeographical factors that shape genetic diversity in western North America.

## Materials and Methods

### Tissue sampling

Tissue samples were collected between June and August of 2018 to 2023 from coastal BC to the eastern edge of the species’ range in Alberta (Fig. 1B; Table S1). Sites across the putative boundary between the Calling and Non-Calling populations were selected to be as close as possible to sites from which individuals were characterized for the presence or absence of vocal sacs (as per Pauly 2008). Between one and five individuals were collected from each site, resulting in a total of 46 tissue samples from 39 sites. Whole larvae (or young of the year when larvae were not available) were captured by hand or by dipnet from each site. Individuals were humanely euthanized on site using a 3% buffered MS-222 solution. Samples were preserved in 95% ethanol before being brought to the lab for long-term storage at -80°C. Fieldwork and sample collection was conducted under permits granted by the Government of Alberta (Alberta No. 22-175 and 23-185), Government of British Columbia Ministry of Forests (MRNA16-236743, MRCB23-787858), Parks Canada (WTNP-2023-45325), and the Solutions Table of the Council of the Haida Nation and the Haida Gwaii Natural Resource District (Gamlen-Greene Haida Gwaii Amphibian Research 03-Nov-16). Samples were collected under approved Canadian Council for Animal Care protocols to different members of the research team (University of Lethbridge protocol #1912, University of Ottawa protocol #3991, Government of British Columbia 236743-ACA, and University of British Columbia protocols A16-0210 and B16-0222).

### DNA extraction and sequencing

Total genomic DNA was extracted from 25 mg of larval tail tissue or metamorphic leg muscle using Qiagen DNeasy Blood and Tissue Kits (Qiagen, Valencia, CA, USA). The manufacturer’s protocol was followed with the following modifications: We added a physical homogenization step, using a VWR bead mill with 2.8 mm ceramic beads to improve tissue digestion. Tissues were then digested for three hours at 56°C, after which, we added 6 μl of RNAase A solution (10 mg/ml). Samples were eluted in two steps, using 80 μl and 60 μl of buffer AE respectively. DNA concentration and quality were assessed using a Thermo Fisher Qubit 4 fluorometer (Thermo Fisher Scientific, Waltham, MA, USA) and NanoDrop 1000 spectrophotometer (Nanodrop Technologies Inc. Wilmington, DE, USA). Samples were diluted and normalized to a concentration of 20 ng/μl in 20 μl (400 ng total) before library preparation.

We obtained sequence data for each individual using double digest restriction site-associated DNA sequencing (ddRADseq; Petersen et al., 2012). Library preparation was done at the Plateforme d’Analyses Génomiques of the Institut de Biologie Intégrative et des Systèmes (IBIS, Université Laval, Quebec, Canada) and was based on the genotyping-by-sequencing procedures described by Poland et al., (2012). Double-stranded DNA was digested using the restriction enzymes *PstI* (5-CTGCAG-3) and *MspI* (5’-CCGG-3’). A set of corresponding Illumina adapters were ligated to the ends of the fragmented DNA and the inline barcodes were ligated to the adapters. Prior to PCR amplification, a Blue Pippin (SAGE Sciences Inc. Beverly, MA, USA) was used to select fragment sizes between 100 and 300 bp using an elution time set between 50 and 65 minutes on a 2% gel. Individuals were sequenced in a single library that included both a technical replicate (replicated from DNA extraction) and two negative controls. A single library was sent to the Centre d’expertise et de service Génome Québec at McGill University in Montreal Quebec for paired-end sequencing on an Illumina NovaSeq 6000. There, multiple libraries were normalized, pooled, and then denatured in 0.02N NaOH and neutralized using HT1 buffer. Using the Xp protocol as per the manufacturer’s recommendations, pools were loaded at 200 pM on an Illumina NovaSeq S4 lane with 2 x 150 reads on a 300-cycle run in paired-end mode. Base calling was performed using RTA v3 and bcl2fastq2 v. 2.20 (Illumina Inc., 2019) was used to demultiplex the libraries. A total of 1,506,075,128 reads were generated across all individuals.

### Loci assembly and filtering

Raw reads were demultiplexed using *process_radtags* in Stacks (version 2.66 Catchen et al., 2013). Removing reads with uncalled bases or adaptor contamination left 971,781,833 reads across all individuals. Fastp (version 0.23.4, Chen, 2023) was used to trim barcodes and adapters by removing five and three bases from the forward and reverse reads, respectively. We also used Fastp to remove reads with an average Phred quality score less than 20, resulting in 929,487,304 reads (1,853,538 to 56,194,548 reads per individual). We then used multiqc (version 1.21, Ewels et al., 2016) to verify sequencing quality and to ensure GC content along the reads was not excessive.

Reads were aligned to the western toad reference genome (Jackson County, Colorado, USA; Trumbo et al., 2023) using the Burrows-Wheeler algorithm in bwa-mem (version 0.7.18, Li and Durbin, 2009). The resulting SAM files were sorted and converted to BAM format using Samtools (version 1.20, Li, 2011). We called single nucleotide polymorphisms (SNPs) from across mapped regions of the reference genome using the *gstacks* module in Stacks and filtered out stacks with a quality score less than 20. SNPs with high heterozygosity often result from paralogs or alignment errors. To remove SNPs with excess heterozygosity, we set the maximum observed heterozygosity (--max-obs-het) to 0.5 in the *populations* module in Stacks, treating all individuals as a single population.

Further filtering was conducted in R (version 4.3.3, R core team, 2024), using the package SNPfiltR (DeRaad, 2023). To ensure accurate genotype calls, we removed SNPs with a genotype depth of less than five or with more than 22 reads. We additionally removed genotype calls with quality of less than 20. The ratio of reads with reference versus alternative bases in heterozygous individuals is expected to be balanced, with large imbalances potentially signaling issues with coverage, multilocus contigs or other artifacts (O’Leary et al., 2018). Thus, we filtered loci with an allele balance less than 0.25 or greater than 0.75 (i.e. the minimum number of reads to call a heterozygote was two). We then applied a minor allele count of three (i.e. with mac = 3, there must be at least one homozygote and one heterozygote for the alternative allele across all individuals to keep a SNP). We iteratively filtered for both missingness of individuals and SNPs (e.g. O’Leary et al. 2018). Given that most sites in this study are represented by a single individual, we prioritized the retention of individuals over SNPs during this process. Finally, we removed SNPs that are likely in linkage disequilibrium using the function *--indep-pairwise* in PLINK (version 1.90b, Chang et al., 2015). This function calculates the correlation coefficient between pairs of SNPs within a specified window and removes SNPs until all the correlations within the window are below a given threshold. We used a window size of 50 base pairs and tested step sizes of 5 or 10 base pairs and R^2^ threshold values between 0.5 and 0.8 before settling on a final step size of 5 and R^2^ of 0.8 (see Supplementary Methods). Additional details of loci assembly and filtering, including the number of SNPs retained at each step, can be found in the Supplementary Methods.

### Characterizing genetic structure across the range

We used four approaches to assess genetic structure across the Canadian range of western toads. First, we visualized the distribution of individuals in multivariate genetic space using principal component analysis (PCA) and discriminant analysis of principal components (DAPC). PCA was run using PLINK (Chang et al., 2015). DAPC was run using the R package Adegenet (Jombart, 2008). Prior to running DAPC, we used the *find.clusters* function to determine the optimal number of genetic clusters based on the Bayesian Information Criterion (BIC) and to assign individuals to these clusters. We retained four principal components and two discriminant functions.

In addition to ordination analyses, we assessed phylogenetic relationships between individuals by generating a maximum likelihood tree in IQTree (version 2.2.2.7, Minh et al., 2020), using the ascertainment bias correction (ASC) to account for the lack of invariant sites. Use of the ASC required removal of partially consistent sites (SNPs for which one variant is ambiguously called). Removing these sites left 9,648 SNPs for this analysis. IQTree was run with 1000 Ultrafast bootstrap replicates using the TMV+F+ASC+G4 model of sequence evolution, which was identified as the best model by ModelFinder (Kalyaanamoorthy et al., 2017).

Finally, we used the admixture model in STRUCTURE (version 2.3.4, Pritchard et al., 2000) to estimate the proportion of each individual’s ancestry derived from inferred genetic groups. STRUCTURE requires the user to specify the number of groups (K) *a priori* and we tested values of K from 1 to 10. Structure was run with a burn-in period of 300,000 steps, 1,000,000 Markov Chain Monte Carlo (MCMC) steps, and 10 replicate runs for each value of K. CLUMPAK (Kopelman et al., 2015) was used to average the runs for each value of K, and ΔK (Evanno et al., 2005) was used to determine the optimal value of K.

All analyses revealed the same clustering of individuals (see Results), and we used the R package dartR (Mijangos et al., 2022) to estimate F_ST_ between genetic groups. To understand the role of IBD in structuring genetic diversity within the species’ range, we used Mantel tests to determine if there was a significant association between pairwise geographic and genetic distance. Because most sites in the dataset were represented by a single individual, we used the proportion of shared alleles between pairs of individuals as an estimate of genetic distance (rather than differences in allele frequencies). To avoid issues of non-independence associated with multiple individuals at some sites, for the few sites with multiple individuals, we randomly selected a single individual for this analysis. Genetic distances between individuals were calculated using the *gl.dist.pop* function from the dartR package and pairwise Euclidean geographic distances between individuals were calculated using the *distm* function in the R package geosphere (Hijmans, 2024a). Separate Mantel tests were run for: 1) all samples, 2) all samples excluding the three individuals from southeastern BC and southwestern Alberta that formed a separate genetic group (see Results), 3) all samples excluding two individuals from island sites in BC, 4) individuals from the Calling range and 5) individuals from the Non-Calling range. Mantel tests were run in R using the *gl.ibd* function from the dartR package.

Although patterns of IBD are expected in widespread species with continuous distributions (Turbek et al., 2023), if there is a sharp genetic break in the range, we expect estimates of pairwise genetic distances between groups to be higher than estimates within groups at all distance classes, including over relatively short geographic distances—that is we expect differences in the y-intercept in a plot of geographic distance versus genetic distance when pairs of samples are evaluated by comparison type. To test for a genetic break in IBD associated with the boundary between the Calling and Non-Calling populations, we focused on pairwise genetic distances between individuals that were separated by no more than 150 km (less than the average width of the CRMs), noting the comparison type in each case (Calling to Non-Calling: CN, Calling to Calling: CC, Non-calling to Non-Calling: NN). We then calculated the difference in mean pairwise genetic distances between the different comparison types (i.e. mean CC versus mean CN; mean NN versus mean CN). To test the statistical significance of these mean differences, we compared the observed mean difference in each case to a null distribution of differences in means generated by randomizing comparison type assignment across the pairwise genetic distance values 1000 times. Mean differences greater than the 95^th^ percentile of this distribution were considered to be statistically significant (i.e. one-sided test of significance).

### Concordance between genetic boundaries and call morphology

Pauly (2008) previously examined 201 museum specimens of sexually mature, male, western toads from 84 sites across the Canadian portion of the range. Individuals were scored for the presence of vocal slits (from which vocal sacs originate). Specimens with one or both vocal slits were scored as Calling whereas those without vocal slits were scored as Non-Calling (Pauly, 2008). We assigned the 39 individuals from that study with accurate GPS coordinate information and that were within 50 km of a genetic sample (a reasonable distance for life-time dispersal in this species: e.g. Bull, 2006; Thompson, 2019) to genetic clade based on Euclidean distance to the nearest genetic sample included in our phylogenetic tree. We then used a Fisher’s exact test to test the association between morphological classification and genetic clade assignment.

### Ecological niche models and niche overlap

Ecological Niche Models (ENMs) have emerged as an important tool in molecular ecology for assessing ecological differences between genetic groups (e.g. Campbell et al., 2022; Gallego-Garcia et al., 2023; MacDonald et al., 2025b) and the potential role of ecological barriers in maintaining groups (e.g. Glor and Warren, 2011; Lee-Yaw and Irwin, 2015). We generated separate ENMs for the Calling and Non-Calling populations to explore the climate variables that best predict presence and the distribution of suitable habitat for each population. Species occurrence records for the models were downloaded from the Global Biodiversity Information Facility (GBIF; December 26, 2024) and filtered to include occurrences from 1990 onward. To match the resolution of the environmental variables (below), we retained records with an accuracy of <1 km and retained one occurrence per grid cell of the environmental variables. Occurrence records were classified as belonging to the Calling (N = 215) or Non-Calling (N = 1523) population based on the recognized morphological boundaries (Pauly 2008; Environment and Climate Change Canada, 2020). Thirty-one climate variables representing climate normals for the period between 1990 and 2020 were downloaded from ClimateNA (Wang et al., 2016; Mahony et al., 2022) at a resolution of 30 arc seconds (1 km by 1 km) and were cropped to the boundaries of Alberta and BC. To avoid issues with collinearity in the models, we removed variables with a high variance inflation factor (VIF >10; calculated in R: usdm Naimi et al. 2014), retaining eight variables for the models: Julian date on which the frost-free period begins, degree-days below 18°C, mean temperature of the coldest month, precipitation as snow (mm), summer (June to August) precipitation (mm), spring (March to May) precipitation (mm), mean annual relative humidity (%), and summer heat moisture index.

Niche models were generated in the R package predicts (Hijmans, 2024b) using Maxent (version 3.4.3, Phillips et al., 2006), which often outperforms other presence-only modeling algorithms (Elith et al., 2011; Bradie and Leung, 2017). Separate models were generated for the Calling and Non-Calling population using the currently recognized range of each population in Canada (Environment and Climate Change Canada, 2020). To reduce the impacts of sampling bias in our models, we sampled 5000 background points for each model from a bias grid corresponding to the density of observations of pond-breeding amphibians in each study extent (i.e. target-group background as per Barber et al., 2022; Baker et al. 2024). Records for the target-group (American bullfrog, boreal chorus frog, Canadian toad, Columbia spotted frog, long-toed salamander, northern leopard frog, northern pacific tree frog, northwestern salamander, western tiger salamander, wood frog) were retrieved from GBIF on October 21, 2021 and were filtered in a manner consistent with the occurrence records (above). The bias grid was generated by applying a Gaussian kernel density estimation to the target-group occurrences using the *density.ppp* function from the spatstat.core package in R (Baddeley et al., 2023). We used the ENMeval package (Kass et al., 2021) in R to select the optimal regularization multiplier (RM) parameter and feature class setting for the Maxent models. Values of RM tested during tuning were: 0.5, 1, 1.5, 2, 2.5, 3, 3.5, 4. Feature classes tested were: L, LQ, H, LQH, LQP, LQT, LQHP, LQPT, and LQHPT where L= linear, Q = quadratic, H = hinge, P= product and T = threshold. The Akaike information criterion correction (AICc) was used to select the final combination of RM and feature classes with which to generate each model. Models were generated with the convergence threshold set to 0.001% (Phillips et al., 2006) and the number of maximum iterations set to 5000 to allow models to converge (Young et al., 2011). All other Maxent settings were kept as default values, including prevalence (𝜏 = 0.5), which has produced accurate models for this species in other contexts (e.g. Bergman et al., 2024). For each model, we determined the area under the receiver operating characteristic curve (AUC) based on average AUC from spatial block (4-fold) cross validation.

Models were projected across BC and Alberta using Maxent’s cloglog output, resulting in a continuous prediction surface for each population. To avoid extrapolation, we used the *mess* function in the R package *predicts* to generate a Multivariate Environmental Similarity Surface (MESS; Elith et al., 2010) for each population and used this surface to mask areas in our prediction surfaces with conditions outside those used to calibrate the models. These areas can be treated as harboring environmental conditions that are “untested” by the population in question. To facilitate interpretation, we converted continuous model predictions into binary surfaces of suitable habitat using the 10% omission rate of the input locality records as a threshold for calling a cell suitable or not (Vale et al., 2014; Rosner-Katz et al., 2020; Bergman et al., 2024). We used both the continuous and binary predictions of suitable climatic space to assess how climate may influence the boundaries of each population and the potential for gene flow between the two populations.

We additionally visualized the amount of niche overlap between the Calling and Non-Calling populations based on all 31 ClimateNA variables using the PCAenv approach of Broennimann et al. (2012) as executed by the *ecospat.plot.niche.dyn* function in the R ecospat package (Broennimann et al., 2025). We used the *ecospat.niche.dyn.index* function in ecospat to calculate niche stability (Guisan et al., 2014). In our case, because both the accessible (background) and occupied environmental space for the Calling population were almost entirely nested within the much larger, occupied environmental space of the Non-Calling population (see Results), we used the test to ask what proportion of the Calling population’s occupied niche is also occupied by the Non-Calling population (i.e. test configured with the Calling population as the “native species” and D restricted to the shared environment space occupied by both populations). Finally, we used the *ecospat.niche.similarity.test* function in ecospat (Warren et al., 2008 and Broennimann et al., 2012) to quantify niche overlap between the two populations using Schoener’s D (Schoener, 1968). D ranges from 0 (no overlap) to 1 (complete overlap) and we tested whether the observed value of D was greater than a null distribution of values generated by fixing the observed niche of one population and randomly selecting a niche for second population within its range 1000 times. This test asks whether the niche of the population being randomized is more similar to that of the other population than expected by chance, given its accessible environment. We ran the test twice, alternating which niche was held constant.

## Results

### Sequencing and filtering

After filtering and removing individuals with large amounts of missing data, our final genomic dataset consisted of 40 individuals from 34 sites distributed from coastal BC to the eastern edge of the species’ range in Alberta and inclusive of populations in and adjacent to the CRMs (Fig. 1B). Individuals were genotyped for 11,950 SNPs. Mean depth of coverage was 19.0X, with no more than 5% missing data per SNP and less than 33% missing data per individual.

### Pronounced genetic structure across the Canadian range of western toads

All analyses assessing genetic structure clearly identified two groups of western toads corresponding to what are referred to as the Calling population to the east of the CRMs and the Non-Calling population to the west of the CRMs (hereafter we will use Calling and Non-Calling populations to refer to both the current Designatable Units and the genetic groups interchangeably). Specifically, Calling and Non-Calling toads separated along PC1, which explained 27.6% of the variation in the principal components analysis (Fig. 2A). Similar separation between the two groups was confirmed by the DAPC analysis (Fig. S1). Calling and Non-Calling individuals also formed distinct clades in the maximum likelihood tree, with bootstrap support values of 100 and 96 respectively (Fig. 2B). Finally, K of 2 in the STRUCTURE analysis received the highest support (ΔK value of 5146; Fig. S2), with Calling and Non-Calling individuals forming separate groups (Fig. 2C). Although there were some admixed individuals along the boundary between groups (Fig. 2D), average assignment of Calling and Non-Calling individuals to their respective groups was high (Q_Non-Calling_ = 0.94; Q_Calling_ = 0.98).

**Figure 2.**
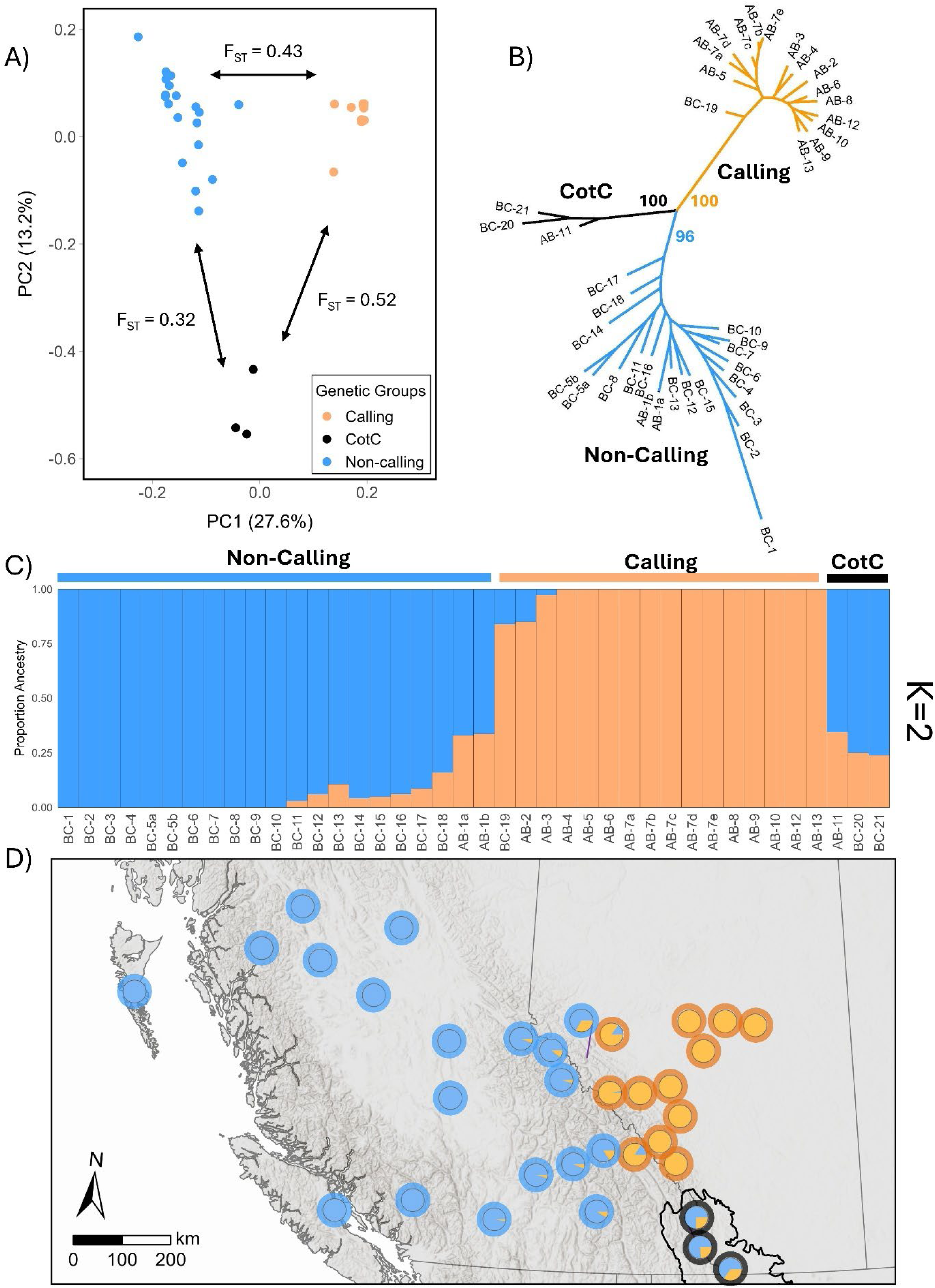
Population structure within western toads (*Anaxyrus boreas)* across the Canadian portion of the species’ range. Principal components analysis revealed three discrete genetic clusters corresponding to Calling and Non-Calling individuals, as well as an additional group (CotC) associated with the Crown of the Continent Ecosystem (A). These groups formed monophyletic clades in the unrooted maximum likelihood tree, with high bootstrap support (B). Two groups (K = 2) were optimal in the STRUCTURE analysis, representing Calling and Non-Calling individuals (C), which are separated by the Canadian Rocky Mountains (D). In all panels, orange symbols and shading represent the Calling population, blue symbols and shading represent the Non-Calling population, and black symbols and shading represent individuals from the CotC. Symbols in D represent locations from which individuals were sampled (one individual per location) and show the proportion of each individual’s ancestry to the two groups identified by the STRUCTURE analysis at K = 2. The colours of the thick borders around all points represent clade membership based on the maximum likelihood phylogeny in B. The extent of the Crown of the Continent Ecosystem in the south is indicated by the black polygon.

In addition to the differences between the Calling and Non-Calling populations, our results revealed a third genetic group in Canada. Specifically, individuals from three sites in southwestern Alberta and southeastern BC separated from the Calling and Non-Calling populations along PC2, which explained 13.2% of the variation in the principal components analysis (Fig. 2A). These sites are located in what is referred to as the Crown of the Continent Ecosystem (UNESCO, 2025; Fig. 2D) and are hereafter referred to as the CotC group. The DAPC analysis with the lowest BIC (value of 231) supported K = 3, with the CotC group also forming a distinct group in this analysis (Fig. S1). Likewise, the CotC individuals formed a separate clade in the phylogenetic tree with bootstrap support of 100 (Fig. 2B). Although these individuals appear admixed in the STRUCTURE analysis when K = 2, they form a separate group at K = 4 (K=3 pulled out the Calling and Non-calling populations and a group for which the biological significance was unclear; Fig S3). Genetic differentiation between all three genetic groups was high, with F_ST_ between the CotC group and the Calling (F_ST_ = 0.52) and Non-Calling populations (F_ST_ = 0.32) similar to or higher than between the Calling and Non-Calling populations (F_ST_ = 0.43).

Across all sites there was a significant pattern of IBD (Mantel R^2^ = 0.60, p < 1.0e-4; Fig. 3A). Significant IBD was also observed when focusing on individuals from either the Calling or Non-Calling populations (i.e. excluding CotC individuals: Mantel R^2^ = 0.61, p < 1.0e-4) and when removing individuals from island sites (Mantel R^2^ = 0.56, p < 1.0e-4). We also found a significant pattern of IBD within the ranges of both the Calling and Non-Calling populations (Calling: Mantel R^2^ =0.36, p < 0.05; Non-Calling: Mantel R 0.69, p < 1.0e-4; Fig. 3B).

**Figure 3.**
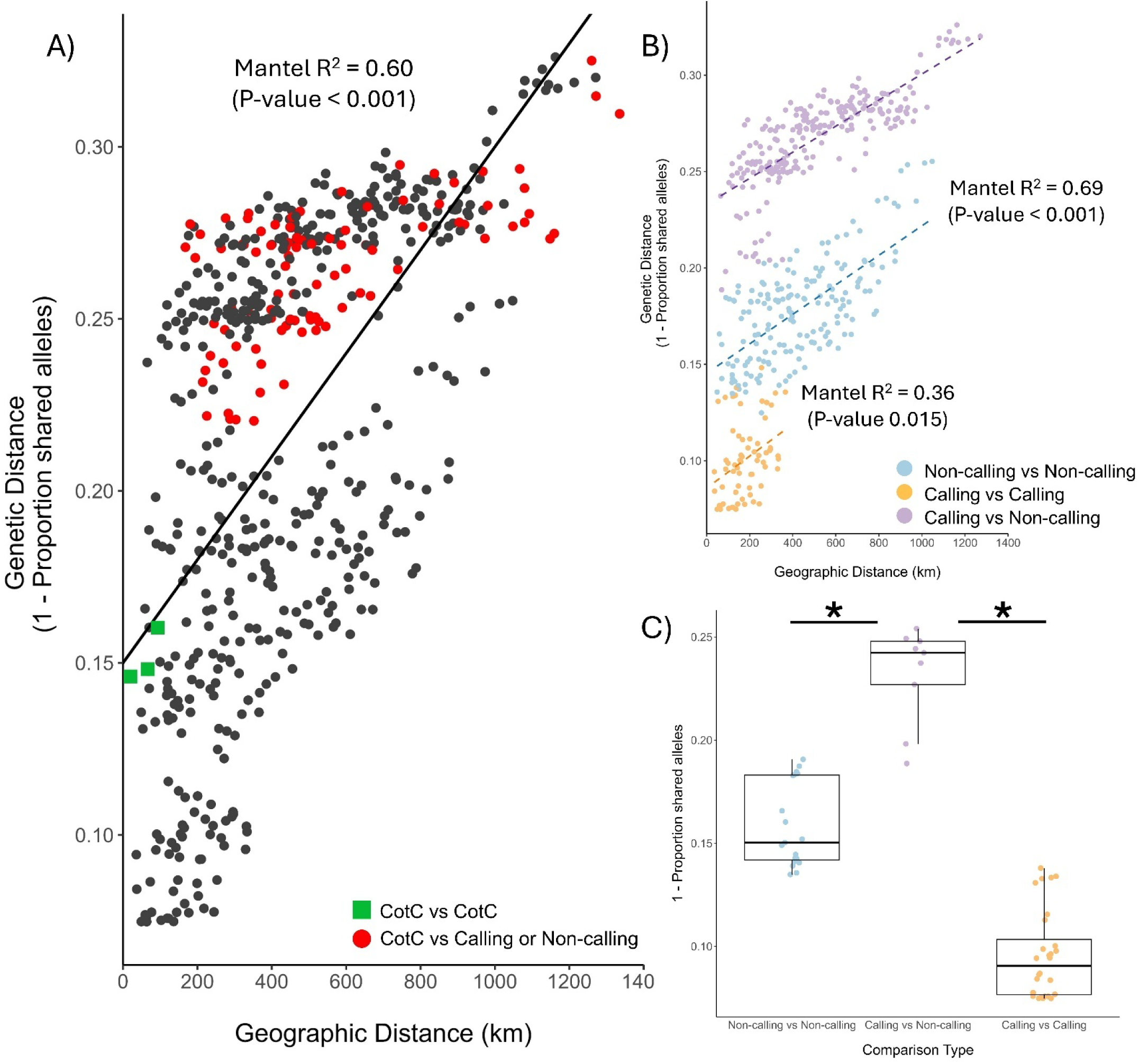
Pairwise comparisons of geographic distance (km) versus genetic distance (1 – proportion of shared alleles) for different subsets of the data (A and B). (A) The solid black line is the Mantel test regression line of the relationship between geographic and genetic distance (IBD) for all individuals. Green squares are pairwise comparisons of “CotC” to “CotC” individuals, and red points are pairwise comparisons between “CotC” individuals and either Calling or Non-Calling individuals. (B) The same plot from A but with comparisons involving the CotC group removed and points color-coded by comparison type for clarity (orange: pairs of individuals within the Calling population; blue: pairs of individuals within the Non-Calling population; purple: pairs of individuals from different populations). The orange and blue dashed lines are the Mantel test regression lines from the isolation-by-distance analyses for the Calling and Non-Calling populations respectively. The regression line through Calling vs Non-Calling comparisons (purple dashed line) is for illustration purposes only. (C) Mean pairwise genetic distances for different comparison types for those individuals within 150 km of each other. Asterisks above the bars connecting between-population comparisons indicate significance based on a randomization test.

We found evidence for a sharp genetic break across the boundary between Calling and Non-Calling individuals. Specifically, across all geographic distances, pairwise genetic distance when comparing Calling to Non-Calling individuals was higher than when comparing Calling to Calling individuals or Non-Calling to Non-Calling individuals (Fig. 3B). Importantly, the mean genetic difference between Calling and Non-Calling individuals was significantly greater than the mean genetic differences between individuals from the same clade over relatively small distances (up to 150 km), including between sites spanning the boundary between genetic groups (mean difference of proportion of shared alleles CC vs CN = 0.137, p-value <0.05; mean difference of proportion of shared alleles NN vs CN = 0.074, p-value <0.05; Fig S4; Fig 3C). There were too few individuals in the third CotC group to test for IBD within this group. However, we note that pairwise genetic distances between the three individuals in the CotC group (green squares: Fig. 3A) were all smaller than pairwise genetic distances between each of these individuals and individuals in either the Calling or Non-Calling populations (red points: Fig. 3A).

### Agreement between genetic boundaries and call morphology

We found a significant association between call morphology and genetic assignment (Fisher’s exact test, p-value = 2.5e-07). All 14 individuals that were morphologically Calling (i.e. presence of one or more vocal slits) were geographically closest to sites that fell into the Calling genetic clade in the phylogenetic tree (Table 1). Of the 25 individuals that were morphologically Non-Calling (i.e. absence of vocal slits), the majority (17) were geographically closest to individuals belonging to the Non-Calling genetic clade (Table 1). Of the remaining eight morphologically Non-Calling individuals, four were from one site, which was geographically closest to an individual in the Calling genetic clade (i.e. mismatch between call morphology and genetic assignment). Notably, this site also included a morphologically Calling individual and was the only mixed phenotype site included in the analysis. This site falls along the genetic boundary between the Calling and Non-Calling groups and is an area of genetic admixture based on the STRUCTURE analysis (Fig. 4). The remaining four morphologically Non-Calling individuals were geographically closest to the CotC genetic group (Table 1).

**Figure 4.**
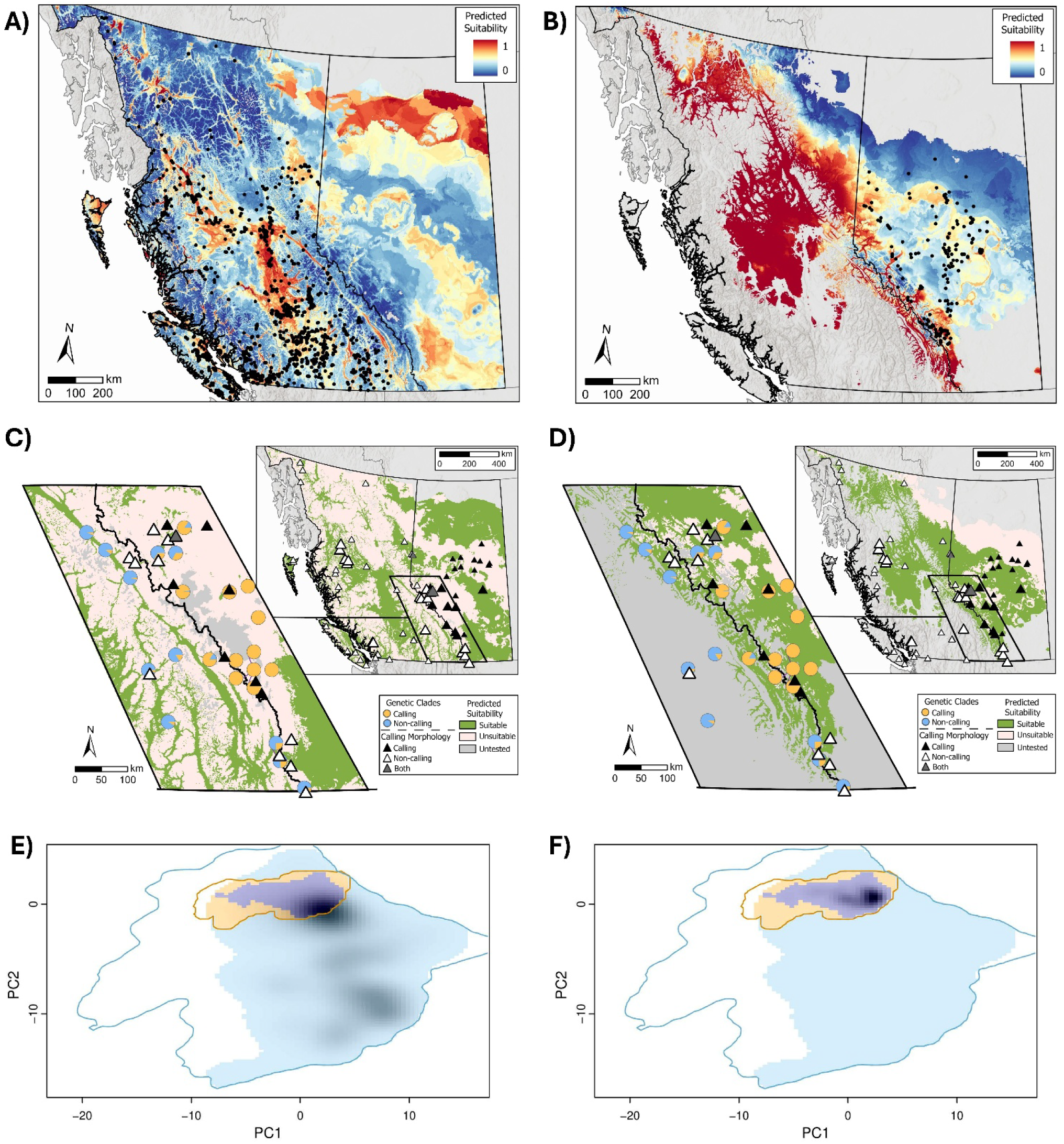
Analysis of niche overlap between Calling and Non-Calling populations of western toads in Canada. Comparison of continuous (A, B) and binary (C, D) predictions of the distribution of suitable habitat for the Calling (left panels) and Non-Calling (right panels) populations based on Maxent models. Black points in A and B represent the input occurrence records for each model. Warmer colors in these panels represent areas of higher predicted suitability; cooler colors represent areas of lower predicted suitability. Binary surfaces (insets in C and D) were generated from the continuous surfaces using the 10% omission threshold based on the corresponding occurrences, with green shading showing suitable habitat and light pink shading showing unsuitable habitat. A zoomed-in, arbitrary 400 x 650 km area around the boundary between genetic groups is shown in C and D to facilitate comparison of suitable habitat towards this boundary. Triangles in C and D show locations where individuals were assessed for call morphology, with black triangles showing sites where all individuals had vocal slits (Calling) and white triangles showing sites where all individuals lacked vocal slits (Non-Calling; data originally presented in Pauly 2008). Gray triangles show site along the boundary between groups where both phenotypes were found in the same location (see main text). Larger triangles represent locations used to test the concordance between genetic and morphological groups. Circles in the zoomed-in region show the genetic sites and the proportion of each individual’s ancestry assigned to the two populations based on the STRUCTURE analysis (see Fig. 2D). In all surfaces, gray areas show places where conditions were outside of the range of conditions used to calibrate the models (i.e. untested climatic space) as identified through a Multivariate Environmental Similarity Surface (MESS) analysis. Panels E and F show the available (polygon outlines) and occupied (shading) climatic space for the Calling (orange) and Non-Calling (blue) populations respectively based on 31 climatic variables. Darker shading within the occupied climatic space represents the density of occurrence records for the Non-Calling (E) and Calling (F) population, with dark blue pixels in both panels showing the overlap between populations.

**Table 1.** The association between call morphology (presence or absence of vocal slits) and genetic group membership based on the geographically closest genetic site. Only morphologically-characterized specimens within 50 km of a genetic sample were used.

|  | *Calling morphology* | *Non-calling morphology* |
| --- | --- | --- |
| *Calling genetic group* | 14 (100%) | 4 (16%) |
| *Non-calling genetic group* | 0 (0%) | 17 (68%) |
| *CotC genetic group* | 0 (0%) | 4 (16%) |

### Distribution of suitable habitat and niche overlap

The Maxent models performed well at predicting presence based on spatial block cross-validation (Non-Calling: AUC = 0.73, Calling: AUC = 0.73; Table S2). Both populations responded similarly to most climatic variables (Fig. S5) and both models suggested that the number of degree-days below 18°C strongly influences the probability of presence (permutation importance 18.4% and 33.7% for Calling and Non-Calling models respectively), with the probability of presence decreasing with a greater number of cooler days in the year for both populations. However, most of the other variables deemed to be important for predicting presence by the models were different for the two populations (Table S3). For example, the variable with the highest percent contribution to the Non-Calling population model was the amount of precipitation as snow (44%), which showed a threshold-like effect in terms of probability of presence for this population, but did not have much of an influence on the probability of presence for the Calling population (Fig. S5). In contrast, the Julian date on which the frost-free period begins had the highest percent contribution to the model for the Calling population (38%) and showed a positive relationship with probability of presence with this population, but did little to explain the probability of presence for the Non-Calling population (Fig. S5).

Areas of high climate suitability for the Non-Calling population were predicted throughout the interior plateau of BC, on Haida Gwaii, Vancouver Island, and in parts of Alberta (Fig. 4A, 4C). Areas of high suitability for the Calling population were predicted in central Alberta, in parts of the CRMs, and in the center of BC (Fig. 4B, 4D). There were parts of both BC and Alberta that were predicted to be suitable for both populations, including an area to the north of our transects along the BC-Alberta border (Fig. 4A and B). However, much of the southern CRMs and the general area associated with the genetic and phenotypic transition between the Calling and Non-Calling populations was predicted to be unsuitable for the Non-Calling population (Fig. 4C) and was outside the range of conditions experienced by the Calling population within its geographic range (i.e. “untested by this population”; Fig. 4D).

Both the available and the occupied climatic space for the Calling population within its range were almost entirely nested within the much broader, climatic space occupied by the Non-Calling population (Fig. 4E, 4F). Within the set of conditions found in the Calling population’s range, almost all conditions occupied by the Calling population were also occupied by the Non-Calling population (niche stability index = 0.89). Schoener’s D was very low (D = 0.005) within the climatic space available to both populations and was not significantly greater than when drawing the Non-Calling population’s niche at random from the much larger set of conditions available to it within its range (p-value = 0.309; Fig. S6). In contrast, D was significantly greater than values obtained by randomizing the niche of the Calling population (p-value = 0.048; Fig. S7)—that is, the niche of the Calling population is significantly concentrated within that of the Non-Calling population within the limited environmental space available to both populations.

## Discussion

We aimed to understand population structure across the Canadian range of western toads in Canada. Our results revealed two genetic groups corresponding to Calling and Non-Calling populations, as well as a third, currently undescribed group of western toads associated with the northern part of the Crown of the Continent Ecosystem. Our analysis of the distribution of variation in call morphology and suitable climate space for the Calling and Non-Calling populations respectively suggests that the genetic boundary between Calling and Non-Calling individuals aligns closely with a sharp transition in a major reproductive trait as well as gaps in suitable climate conditions that may limit the extent to which these groups interact, at least along southern parts of the CRMs. This work is timely as the status of western toads in Canada, including whether the two western toad populations should continue to be recognized as legal units (i.e. Designatable Units; DUs), is due for re-assessment by the Committee on the Status of Endangered Wildlife in Canada (COSEWIC; species are reassessed ∼10 years and the species was last assessed in 2012). We thus discuss our results in light of the specific criteria used by COSEWIC to guide these decisions.

### Support for the continued recognition of two populations of western toads in Canada

In Canada, Designatable Units (DUs) are established by the Committee on the Status of Endangered Wildlife in Canada (COSEWIC) using two main criteria. The first criterion focusses on the “discreteness” of populations (COSEWIC, 2023). Populations are considered to be discrete if there are heritable traits or markers that distinguish them and/or there are natural barriers to the transmission of heritable traits (i.e. barriers to gene flow: COSEWIC, 2023). Several of the results presented here speak to the discreteness of the Calling and Non-Calling populations. Our genomic results suggest that Calling and Non-Calling populations are distinct across much of their genome, with levels of genetic differentiation similar to, or even higher than, those observed among conservation units within other species (e.g. Moore et al., 2014; Jorge et al., 2022; Gallego-Garcia et al., 2023; MacDonald et al., 2025b) and even among what are considered distinct species (e.g. Baumsteiger et al., 2017; Campbell et al., 2022). These genetic differences are associated with differences in terms of the presence of a pronounced advertisement call, which may influence conspecific attraction (e.g. Buxton et al., 2015) and thus opportunities for the transmission of genetic information. Finally, although our niche models and assessment of niche overlap suggest that the two populations have similar responses to variation in climatic conditions and that the occupied niche of the Calling population is entirely nested within that of the Non-Calling population, much of the area in and immediately adjacent to the CRMs is predicted to be unsuitable for the Non-Calling population and is outside the range of conditions experienced by the Calling population within its range. Along with the limited amount of genetic admixture observed in this region, these results suggest that the CRMs may act as at least a partial barrier to gene flow between the two populations. Together, these results support the continued recognition of the Calling and Non-Calling populations of western toads as discrete groups.

Isolation by distance (IBD) is expected in widespread species that are continuously distributed across the landscape (Wright, 1943; Turbek et al., 2023) and one challenge when assessing the discreteness of potential groups in widespread species is to ensure that such groups are not simply an artifact of IBD (Chambers and Hillis, 2020; MacGuigan et al., 2022; Turbek et al., 2023). For this reason, we sampled individuals from the west coast, through to the eastern edge of the species’ range, ensuring that individuals on either side of the putative boundary between the groups were included. Although we observed strong IBD across the full extent of the study area and within the ranges of each of the populations, IBD alone is unlikely to account for the observed genetic structure. Specifically, genetic distances are larger on average between Calling and Non-Calling individuals than between individuals within either population at each distance class. Importantly, at relatively small geographic distances (<150 km), including between adjacent samples in and around the transition between groups, we found that mean pairwise genetic distance was significantly higher than expected when comparing samples across populations than when comparing samples within populations. Thus, while IBD plays a role in shaping patterns of genetic variation in western toads, Calling and Non-Calling populations show genetic differences above and beyond what is expected to arise from IBD alone.

The second criteria used to delineate DUs in Canada focuses on the evolutionary significance of populations (COSEWIC, 2023). This criterion often emphasizes the distinct evolutionary histories of populations (COSEWIC, 2023). Our focus on the Canadian portion of the species’ range means our understanding of the biogeographic history of western toads remains incomplete. However, it is likely that western toads, like other species (e.g. Hoffman and Blouin, 2004; Klütsch et al., 2012; Cheng et al., 2014), have undergone periods of isolation in separate refugia during the Pleistocene glaciations. Goebel et al. (2009) previously suggested that western toads may have maintained refugial populations in the Klamath-Siskiyou Mountains, near the border of Oregon and California, and expanded north along coastal BC and Alaska.

More recently, Gamlen-Greene (2022) hypothesized that Haida Gwaii may have potentially served as an additional western refugia for the species (Gamlen-Greene, 2022). However, neither study included sites from further east in the species’ range in Canada. The eastern half of the western toad’s range overlaps with other species in eastern BC and Alberta that are thought to have persisted in ice-free areas of the Rocky Mountains (reviewed by Brunsfeld et al., 2001; Shafer et al., 2010) and interior of Idaho (e.g. Lait et al., 2012) during the Pleistocene glaciations, as well as in cryptic refugia within and between the Laurentide and Cordilleran ice sheets in Canada (Shafer et al., 2010). Thus, it seems likely that western toads had access to multiple glacial refugia and that periods of isolation among different populations have shaped the observed genetic structure within the species’ range.

Differences in adaptive traits are also considered by COSEWIC when evaluating the evolutionary significance of different groups (i.e. “DU-wide differences in adaptive traits not found elsewhere in Canada”: COSEWIC, 2023). The presence of a male vocal sac and pronounced advertisement call in a geographically-discrete part of the western toad’s range and absence of these traits elsewhere is highly unique amongst anurans (Pauly, 2008; Elias-Costa and Faivovich, 2025). The adaptive significance of these differences in calling in western toads is unclear, but in other anurans, variation in the amount of calling that males do and the characteristics of their calls (e.g. frequency) is associated with differences in predation pressure (e.g. Dapper et al., 2010; Bonachea and Ryan, 2011) and the acoustic environment (e.g. Goutte et al., 2018; Zhao et al., 2021). Furthermore, differences in call behaviour and characteristics have promoted divergence in some anuran systems (Pröhl, 2003; Funk et al., 2009) and are thought to have driven speciation in some cases (Boul et al., 2007). Future work is needed to assess whether the differences in call behaviour of Calling and Non-Calling populations of western toads are adaptive and/or contribute to reproductive isolation between the populations. However, DU-wide differences in a key reproductive trait make it likely that Calling and Non-Calling populations are on separate evolutionary trajectories.

### Broader implications for the suturing of genetic diversity in western Canada

The places where different species, or populations within species, meet and interact (i.e. “suture zones”) has long captured the interests of biogeographers and evolutionary ecologists (Remington, 1968; Swenson and Howard, 2004; Swenson and Howard, 2005; Wait and Peñalba, 2025). The CRMs are a major geographic barrier that has partitioned biodiversity at multiple scales (Malanson et al., 2018). For instance, the CRMs represent the transition between three ecozones: the Montane Cordillera in BC and the Boreal foothills and Prairies in Alberta. This transition defines the boundaries between different faunal provinces (COSEWIC, 2023) and is where the western range limits of many species meet the eastern range limits of close congeners. The CRMs are also associated with genetic breaks in several taxa that maintain populations on both sides of the mountains (e.g. silky pocket mouse, North American red squirrel, little brown myotis: Jensen et al., 2024; Canadian Lynx: Watt et al., 2021; American goldfinch: Cloutier et al., 2024; Black-capped chickadees, Boreal chickadees: Lait et al., 2024). Our study adds the first amphibian species to this list, further suggesting that the CRMs are an important suture zone in the north.

Our results also revealed a third genetic group located in southeastern BC and southwestern Alberta. A recent survey of genetic differences across the range of western toads in the United States similarly found a distinct genetic group associated with the Rocky Mountains in Glacier National Park, Montana (Colorado US Fish and Wildlife, 2017), which is also in the Crown of the Continent Ecosystem. Although that study did not include samples in Canada, the close geographic proximity of that group to the samples included in this study, makes it likely that these patterns are reflective of the same genetic substructure. Toads in this part of the Rocky Mountains may be unique with respect to their calling behaviour. Although the specimens examined by Pauly (2008) lacked vocal sacs (both those in Canada and in a broader survey which included sites in Montana), there are reports of toads in southeastern BC and southwestern Alberta producing consistent calls (per. communication L-A. Isaac, BC Ministry of Water, Land and Resource Stewardship; Waterton Lakes National Parks Ecological Monitoring program, Parks Canada Waterton Lakes Field Unit 2021), and Pauly (2008) reported males producing quieter, long, pulsed calls at a single location in northern Montana despite the absence of vocal sacs. Formal analysis of recorded calls suggested these vocalizations differ from the calls of individuals in Alberta and may be an “extra-long release call” (Pauly, 2008) but additional acoustic and behavioural analyses are needed to clarify the nature and function of vocalizations in this part of the species’ range.

The CotC group highlights the potential for the Crown of the Continent Ecosystem to harbour unique genotypes. The Crown of the Continent Ecosystem, spanning mountainous regions of the southwest corner of Alberta, southeast corner of BC, and northern Montana, is characterized by large elevational differences, includes a range of different ecosystems from alpine forests to prairie grasslands, and is one of the most biologically diverse places in North America (Prato and Fagre, 2010). This region is thought to have harbored ice-free areas during the last glacial maximum and may have been an important northern refugia for western species (Brunsfeld et al., 2001; Shafer et al., 2010). Although studies specifically testing the phylogeographic importance of this region are lacking, genetic differences between populations in the Crown of the Continent Ecosystem and elsewhere have been reported in other groups (e.g. Long-toed salamanders: Lee-Yaw and Irwin, 2012; Half-moon hairstreak butterflies: Macdonald et al., 2025b). Additional studies focusing on this region would enhance our understanding of the extent to which the Crown of the Continent Ecosystem is a hotspot of genetic diversity.

### Conclusions and Future Directions

This study is one of the first studies to evaluate genomic differences in a widespread amphibian species in western Canada. We found evidence to support distinct genetic groups associated with Calling and Non-Calling populations of western toads in Canada. Paired with phenotypic and ecological differences between these populations, the continued recognition of these populations as separate conservation units (i.e. DUs) seems warranted. At the same time, we note that we did not have samples from the most northern parts of the species range.

Importantly, our niche models revealed areas of high climatic suitability for both populations to the east of the CRMs in areas north of our sampling. Although there are gaps in available morphological information from this part of the species’ range, there is at least one known location in the Grande Prairie region of Alberta where both phenotypes are found (Fig. 4 C,D; Pauly 2008). At this latitude, the CRMs are less steep, valleys are wider, and the foothills extend further east than in southern parts of the CRMs. Thus, it’s possible that there are more opportunities for contact between the two populations in this region. Additional samples from this region and from across more northern parts of the species’ range are thus necessary to fully characterize the genetic boundaries between populations.

An outstanding question at this point is the extent to which the different genetic groups observed are reproductively isolated. Fine-scale sampling across the boundaries between groups to quantify levels of introgression and hybridization, paired with tests of the role of differences in calling behaviour in mate recognition would shed light on this question. Comparison of levels of reproductive isolation on either side of the southern CRMs to levels in those northern areas where climatic suitability is predicted to be high for both the Calling and Non-Calling populations would be particularly interesting, providing the ultimate test of whether any barriers to gene flow established along putatively different colonization routes around a geographic barrier are maintained upon secondary contact (i.e. akin to Moritz et al., 1992; Helbig, 2005). Regardless of whether Calling and Non-Calling populations are reproductively isolated, the presence of a highly distinct genetic group that is geographically contained entirely within Canada (i.e. the Calling population) is an unusual situation (see MacDonald et al. 2025a for a similarly “curious” example). From the perspective of both understanding and managing genetic diversity, our results thus highlight the importance of filling gaps in our assessments of intraspecific genetic variation in previously glaciated areas.

## Acknowledgements

The Western Toads included in this study are found on the traditional Lands of many First Nations and we recognize the ongoing stewardship of the Indigenous Peoples who have cared for this species and their habitats. We thank the following for their support of the field efforts and sample collection: S. Marks, H. Meikle, P. Monterio, B. Cadsand, E. Wind, M. Thomspon, G. Skinner, G. Dare, C. Gill, K. Pearson, T. Einfeldt, and C. Bergeron. We wish to acknowledge C. Cullingham and J. Martin for comments on an earlier thesis version of this manuscript, and N. Kester for discussion on the niche overlap analysis.

## Data Accessibility

All code is available in an online GitHub repository (https://github.com/Jaynabergman/WETO_ddRAD)

Raw reads from all samples will be available upon publication on the Short Read Archive except for the sample from Haida Gwaii. Data from the Haida Gwaii sample are available on request from the BC Ministry of Forests, subject to review by the Council of the Haida Nation.

## Author Contributions

**Jayna Bergman**: Data curation, Formal analysis, Validation, Software, Visualization, Writing – Original Draft. **Juan Enciso-Romero**: Software, Formal analysis. **Gregory B. Pauly**: Data curation, Writing – Review & Editing. **Roseanna Gamlen-Greene**: Data curation, Writing – Review & Editing. **Melissa Todd**: Data curation, Writing – Review & Editing. **Julie A. Lee-Yaw**: Conceptualization, Methodology, Project administration, Funding acquisition, Supervision, Writing – Review & Editing

## Data Availability

All code is available in an online GitHub repository (https://github.com/Jaynabergman/WETO_ddRAD)

