## Supplemental Methods for "Pronounced genetic structure associated with differences in a reproductive trait and climatic barriers in Canadian populations of the western toad (*Anaxyrus boreas*)"

**Supplementary Methods**

All code can be found on the GitHub repository: <https://github.com/Jaynabergman/WETO_ddRAD>

**Summary of Sampling and sequencing**

- **Number of libraries**: 1 library
- **Number of samples per library:** 46 samples plus one technical replicate (including repeat DNA extraction) and two negative controls
- **Final number of individuals included in the study:** 40 individuals
- **Number of sites:** 34 sites
- **Number of individuals per site:** 1 to 5 individuals per site (two sites with 2 individuals, one site with 5 individuals, 31 sites with 1 individual)

**Demultiplexing, trimming, and adaptor removal**

Raw reads (N = 1,506,075,128) were demultiplexed using *process_radtags* in Stacks (V2.66; Catchen et al., 2013) with the adapter sequences defined (--adapter-1 AGATCGGAAGAG --adapter-2 AGATCGGAAGAG). This resulted in removing 525,075,960 (34.9%) reads that contained adapter sequences. After demultiplexing 971,781,833 reads were retained.

We used Fastp (version 0.23.4; Chen, 2023) and Multiqc (version 1.21; Ewels et al., 2016) to make trimming decisions. We specifically aggregated the json files from Fastp in Multiqc without defining any settings and examined the “GC content plot” to determine the number of bases to trim from the forward and reverse reads (Fig. SM-1). Based on these plots we set the -f (read1) and -F (read 2) to 5 and 3 respectively. We also used Fastp to filter for read quality, setting the Phred score to 20 (-q 20). We retained 929,487,304 reads after demultiplexing, removing reads with adaptors, and trimming in this way.

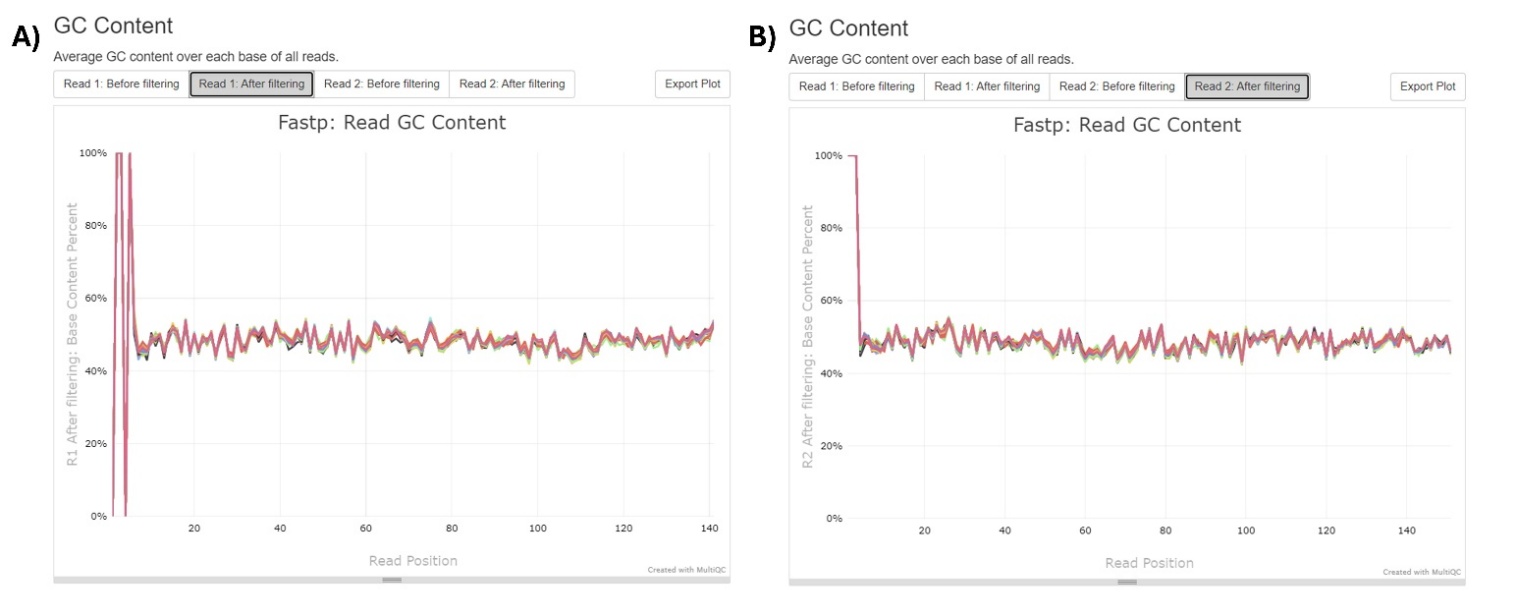

**Figure SM-1**. The average GC content across each nucleotide base of the reads. Each line represents a single individual. These plots were used to determine how many base pairs to trim from the start of the reads for both the (A) forward and (B) reverse reads. Reads were demultiplexed previous to this step and were not otherwise filtered prior to this step.

**Aligning reads to the reference genome using BWA**

The western toad reference genome is approximately 5.2 GB and was developed from a single individual collected in Jackson County, Colorado, USA (Trumbo et al., 2023), and is thus expected to be a member of the Non-Calling clade (i.e. based on Pauly, 2008). This reference genome is comprised of 4709 scaffolds and was sent to us directly by the authors.

We first indexed the reference genome using the program BWA (version 0.7.18). We then used the bwa-mem algorithm to align the reads to the reference genome. SAMtools was used to sort the SAM files and convert them into BAM format. Default parameters were used for the alignment of the reads to the reference genome. A total of 917,788,446 reads uniquely aligned to the reference genome (1,835,041 to 55,541,803 reads per individual).

**Build loci in gstacks (STACKS)**

After aligning to the reference genome, we used the *gstacks* module in STACKS to assemble loci (Table SM-1). *gstacks* identifies SNPs for each locus and then genotypes each individual at these SNPs. To be included at this step, reads had to have a minimum mapping quality (Phred score) of 20 (--minmapq 20).

**Table SM-1.** Output from *gstacks*.

| **Summary Variable** | **Value** |
| --- | --- |
| Number of matching paired end reads | 212,670,599 |
| Read mean insert length | 206.1 (sd: 46.3) |
| Number of genotyped loci | 1,772,249 |
| Mean effective per-sample coverage | 12.0X (ranged from 3.0X to 31.3X, sd: 6.2X) |
| Mean number of sites per locus | 182.3 |

**Filtering SNPs**

We used recommendations from the literature, diagnostic plots, and sensitivity tests to set different filtering parameters. In cases where sensitivity tests were used, we were primarily interested in how filtering decisions might impact conclusions about population structure. As a simple diagnostic in this regard, we relied on principal components analysis (PCA) to assess the stability of the observed genetic clusters to different filtering decisions. Our procedure was as follows:

The *populations* module in STACKS was used to convert individual genotypes to vcf format and to apply a maximum observed heterozygosity filter (--max-obs-het). We tested the sensitivity of results to maximum observed heterozygosity values of 0.5, 0.6, and 0.8. For each iteration, after setting maximum observed heterozygosity, we proceeded to filter SNPs using a reasonable set of starting parameters (minimum genotype depth: 5, genotype quality: 20, allele balance: between 0.25 and 0.75, maximum depth per SNP: 28, minor allele count: 3, and missingness by SNP and missingness by sample iteratively filtered using values of 0.75, 0.85, 0.9 and 0.8, 0.6, 0.4 respectively). Although applying maximum observed heterozygosity values of 0.5, 0.6, and 0.8 resulted in differences in the number of polymorphic sites removed (71000, 56377, and 29884 respectively), after applying these downstream filters, there were no differences in the number or identity of SNPs retained across the different settings of maximum observed heterozygosity. A total of 11,185 SNPs were retained in each case and the resulting PCA plots were identical (Fig. SM-2). We set --max-obs-het to 0.5 for the final filtering scheme.

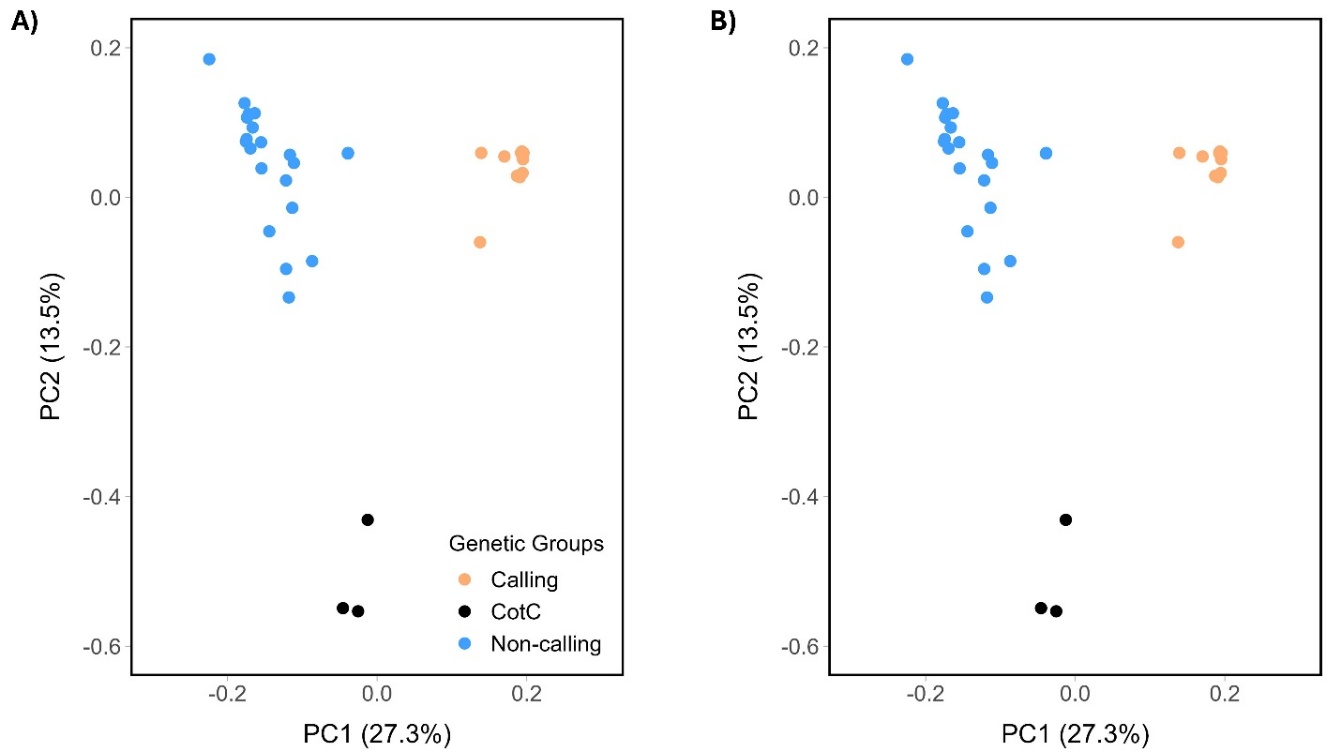

**Figure SM-2**. Varying maximum observed heterozygosity (--max-obs-het) in the *Populations* module in STACKS resulted in an identical set of SNPs being retained once downstream filters were applied and thus did not impact the clustering of individuals to groups in a principal components analysis. Results for values of (A) 0.6 and (B) 0.8 are shown and are identical to results for 0.5, which is shown in the main text.

Additional filtering steps took place in R (version 4.3.3, R core team, 2024) and used the package SNPfiltR (DeRaad, 2023). We used diagnostic plots to set the minimum genotype depth to 5 and the maximum depth per SNP to 22 (Fig. SM-3). Following recommendations from the literature (O’Leary et al., 2018; Hemstrom et al., 2023), we set genotype quality to 20, the allele balance to be between 0.25 and 0.75, and applied a minor allele count (MAC) of 3.

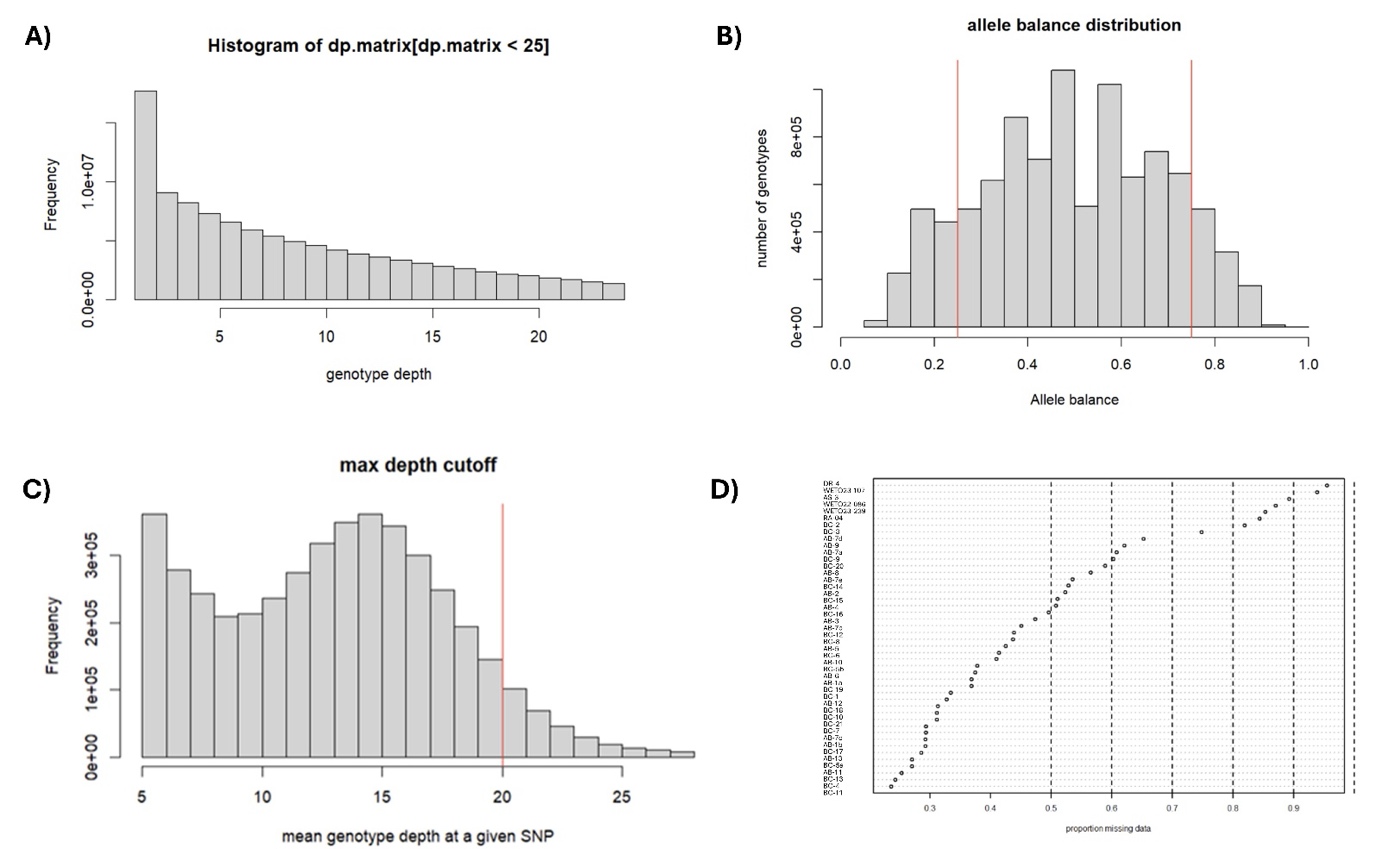

**Figure SM-3**. Graphs used make filtering decisions. (A) Distribution of mean depth per genotype. (B) Allele balance in heterozygous genotypes with red lines indicating standard cut off values of 0.25 and 0.75. (C) Distribution of mean depth per SNP with the selected cutoff of 22 shown as a red line. (D) The proportion of missing data per individual for all 46 individuals before individuals or loci were removed.

We used the statistical approach in the software PLINK to remove SNPs in linkage disequilibrium. This approach requires setting a window size, a step size, and r^2^ cut-off. We used a standard window size of 50 and tested select combinations of common step sizes (5 or 10) with different r^2^ values (0.5, 0.7, and 0.8). The number of SNPs retained ranged from 7,865 to 11,213 depending on the combination of values used (Table SM-2). Step size did not influence the number of SNPs retained as much the r^2^ (Table SM-2). We thus chose the common (and, in this case, slightly more conservative) step size of 5. Using this step size, PCA revealed no major differences in the clustering of individuals into groups when varying r^2^ (Figure. SM-4). Thus, to maximize the number of SNPs retained, we set r^2^ to 0.8.

**Table SM-2**. Different combinations of linkage disequilibrium settings (window, step and r^2^) using the --indep-pairwise function in PLINK, and the corresponding number of SNPs that were removed and kept for each combination. The starting dataset had 20,369 SNPs.

| **Settings (window, step, r2)** | **SNPs removed** | **SNPs kept** |
| --- | --- | --- |
| 50, 5, 0.5 | 12,504 | 7,865 |
| 50, 10, 0.5 | 12,441 | 7,928 |
| 50, 10, 0.7 | 10,235 | 10,134 |
| 50, 5, 0.8 | 9,184 | 11,185 |
| 50, 10, 0.8 | 9,156 | 11,213 |

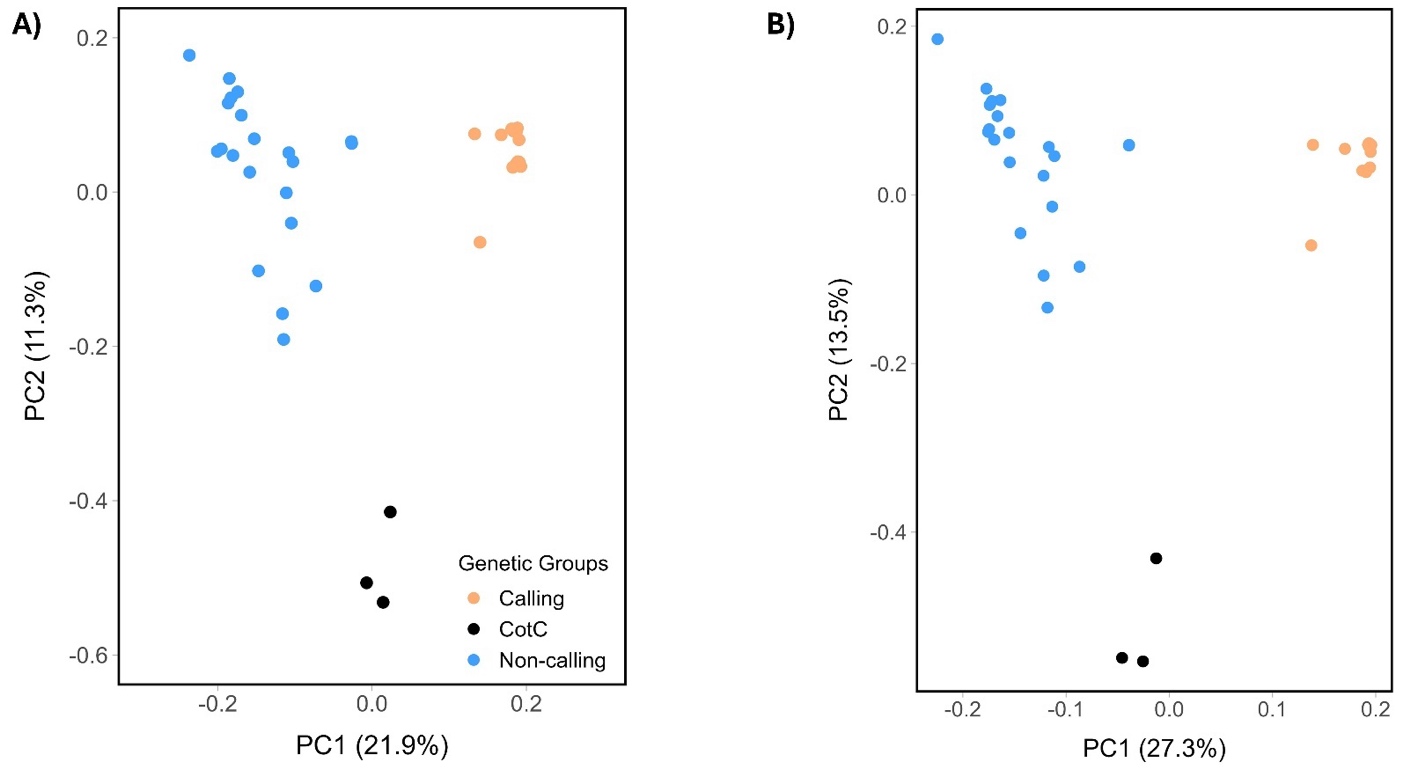

**Figure SM-4**. Setting a more stringent value of r^2^ (i.e. 0.5) to identify SNPs in linkage disequilibrium in the program PLINK did not alter the clustering of individuals into genetic groups. Shown are results from a principal components analysis based on SNPs retained when setting r^2^ to 0.5 (A) and 0.8 (B). In both cases, the window was set to 50 base pairs and the step between windows was set to 5 base pairs. Using an intermediate r^2^ value of 0.7 resulted in qualitatively similar plots (not shown).

Before applying the final LD filter to our dataset, we explored the impacts of the amount of missing data permitted per individual and per SNP the clustering of individuals into genetic group. We started with assessing the amount of missing data per sample and per SNP without removing anything (e.g. Figure SM-3D). We generated a dataset that was more stringent (i.e. less missing data; Figure SM-5A), and others that were more relaxed and thus kept more individuals (i.e. more missing data; Figure SM-5B). Regardless the amount of missingness permitted, we observed similar clustering of individuals in the PCAs (Figure SM-5). To keep as many individuals as possible, we selected a final filtering scheme with the following iterations: 1) remove individuals with missing data >90%, 2) remove SNPs with >85% missing data, 3) remove individuals with missing data >50%, 4) remove SNPs with >95% missing data, and finally 4) remove individuals with missing data >33% (Table SM-3). This resulted in the final dataset to have up to 5% missing data per SNP and up to 33% missing data per individual. We note that there were no clear patterns in the amount of missing data per individual with respect to the genetic groups that emerged from our analyses (Figure SM-6). Table SM-3 outlines the full details of the final filtering scheme.

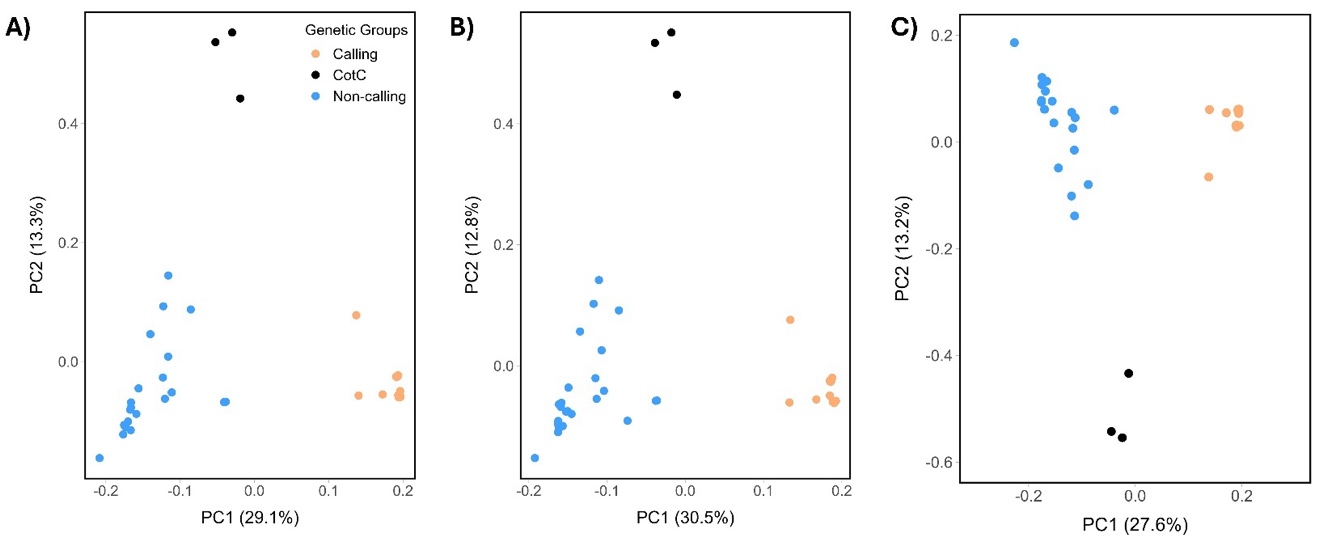

**Figure SM-5**. Principal component analysis (PCA) showing the clustering of individuals into genetic groups when the amount of missingness allowed was varied. (A) Up to 20% missing data per individual and 4% missing data per SNPs (5,031 SNP and 40 individuals retained). (B) Up to 55% missing data per individual and 10% missing data per SNP (10,342 SNPs and 44 individuals retained). (C) Up to 33% missing data per individuals and 5% missing data per SNP (11,950 SNPs and 40 individuals retained). All filtering steps prior to applying these missingness values were identical. The dataset with missingness corresponding to C was used for the final analyses conducted in this study.

**Table SM-3.** Full details of the filtering steps and decisions made to get the final set of 11,950 SNPs used in the downstream analyses.

| **Step** | **Filter/Program** | **How it was decided** | **Setting** | **Reported values** |
| --- | --- | --- | --- | --- |
| 1. | Maximum observed heterozygosity  (--max-obs-het)  In *Populations* (STACKS) | Sensitivity test: Tested multiple values (*see above section; Figure SM-1*). | 0.5 | 71,000 sites filtered out – starting with 4,606,447 variant sites |
| 2. | Genotype depth  In SNPfiltR (R package) | Diagnostic plot (Figure SM-3A): Histogram of the genotype depth frequency; final value is within range recommended in the literature (e.g. O’Leary et al., 2018) | 5 | 28.98% of genotypes fall below a read depth of 5 (converted to NA) |
| 2. | Genotype quality  In SNPfiltR (R package) | Literature standard (O’Leary et al., 2018) | 20 | 0.8% of genotypes fall below a genotype quality of 20 (converted to NA) |
| 3. | Allele balance  In SNPfiltR (R package) | Literature standard (O’Leary et al., 2018; Figure SM-3B); given the specifics of our dataset, heterozygotes to be supported by at least two reads | 0.25 and 0.75  (min and max) | 20.85% of het genotypes [2.2% of all genotypes] fall outside of 0.25-0.75 allele balance ratio (and converted to NA) |
| 4. | Maximum depth per SNP  In SNPfiltR (R package) | Diagnostic plot: Histogram of max depth at a given SNP (Figure SM-3C) | 22 | 4.19% of SNPs removed - 4,410,477 SNPs retained |
| 5. | Minor allele count (MAC)  In SNPfiltR (R package) | Literature standard (Schmidt et al., 2023); given the specifics of our dataset, mac of 3 means allele must be supported by at least one homozygote and one heterozygote | 3 | 50.76% of SNPs fell below a mac of 3 – 2,171,753 SNPs retained |
| 6. | Missing by sample  (missing_by_sample()) in SNPfiltR (R package) | Diagnostic plot (Figure SM-3D): Looked at the plot of missing data by sample before setting any values – two individuals have lots of missing data (>90%), thus we set the filter to 0.9 to remove them. | 0.9 | Two individuals removed (DR4 and WETO23-107) |
| 7. | Missing by SNP  (missing_by_snp()) in SNPfiltR (R package) | Diagnostic table: Looked at missingness by SNP after removing the two individuals above. Used the table that shows how many SNPs are left after removing SNPs with a given percent of missing data. | 0.85 | 96.93% of SNPs fell below a completeness cutoff of 0.85 – 66,625 SNPs retained |
| 8. | Missing by sample | Diagnostic plot: Four individuals are missing >50% of the data. | 0.5 | Four individuals removed (AS-3, WETO22-086, WETO23-239, RA-04) |
| 9. | Missing by SNP | Diagnostic table: Tried to remove SNPs with more missing data in order to keep more individuals | 0.95 | 66.8% of SNPs fell below a completeness cutoff of 0.95 - **22,118** SNPs retained |
| 10. | Missing by sample | Diagnostic plot (Figure SM-6): All individuals had less than 33% missing data. Set this value to confirm. | 0.33 | All individuals have less than 33% missing data (38 individuals have less than 15% missing data) |
| 11. | Linkage disequilibrium | Sensitivity Test: *see section above; Figure SM-4*) | 50, 5, 0.8 | Removed 10,168 SNPs – 11,950 SNPs retained |
| **Final Number of SNPs** | | | | **11,950** |
| **Final Number of Individuals** | | | | **40** |

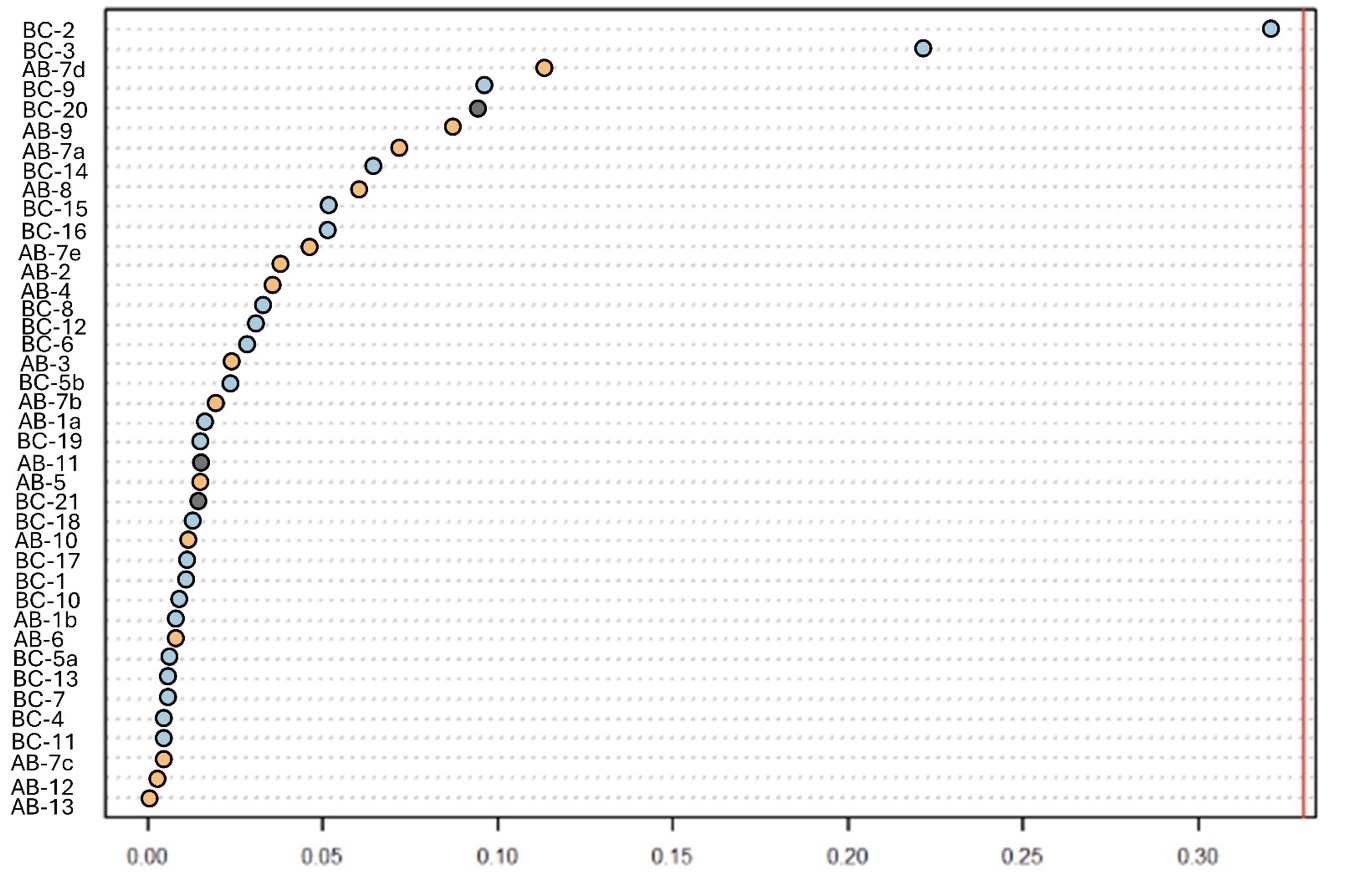

**Figure SM-6**. Final amount of missing data for each of the 40 individuals included in the dataset color-coded by the three genetic groups discussed in the main text (Calling population: orange, Non-calling population: blue; Crown of the Continent group: black).
