## Supplemental Figures and Tables for "Pronounced genetic structure associated with differences in a reproductive trait and climatic barriers in Canadian populations of the western toad (*Anaxyrus boreas*)"

**Table S1**. Tissue sample information for the 40 individuals and the corresponding sites that are included in this study.

| Sample ID | Site ID | Longitude | Latitude | Genetic Clade | Nearest town/county |
| --- | --- | --- | --- | --- | --- |
| BC-1 | BC-1 | Permission required to access | Permission required to access | Non-Calling | Haida Gwaii |
| BC-2 | BC-2 | -128.54 | 54.36 | Non-Calling | Lakelse Lake Area |
| BC-3 | BC-3 | -127.51 | 55.26 | Non-Calling | Ross Lake Provincial Park |
| BC-4 | BC-4 | -126.66 | 54.33 | Non-Calling | Houston County |
| BC-5a | BC-5 | -125.11 | 49.77 | Non-Calling | Vancouver Island |
| BC-5b | BC-5 | -125.11 | 49.77 | Non-Calling | Vancouver Island |
| BC-6 | BC-6 | -124.84 | 53.84 | Non-Calling | Area south of Fraser Lake |
| BC-7 | BC-7 | -124.28 | 55.15 | Non-Calling | Area east of Mudzenchoot Provincial Park |
| BC-8 | BC-8 | -122.94 | 50.13 | Non-Calling | Whistler |
| BC-9 | BC-9 | -122.36 | 53.15 | Non-Calling | Area north of Ten Mile Lake |
| BC-10 | BC-10 | -122.15 | 52.10 | Non-Calling | Williams Lake |
| BC-11 | BC-11 | -120.56 | 49.91 | Non-Calling | Aspen Grove |
| BC-12 | BC-12 | -120.17 | 53.31 | Non-Calling | McBride |
| BC-13 | BC-13 | -119.61 | 53.03 | Non-Calling | Shere |
| BC-14 | BC-14 | -119.45 | 50.79 | Non-Calling | Salmon Arm |
| BC-15 | BC-15 | -118.86 | 52.59 | Non-Calling | Kinbasket Valley |
| BC-16 | BC-16 | -118.24 | 51.03 | Non-Calling | Revelstoke |
| AB-1a | AB-1 | -118.12 | 53.03 | Non-Calling | Jasper National Park |
| AB-1b | AB-1 | -118.12 | 53.03 | Non-Calling | Jasper National Park |
| BC-17 | BC-17 | -117.65 | 50.16 | Non-Calling | Summit Lake Ski and Snowboard Area |
| BC-18 | BC-18 | -117.52 | 51.30 | Non-Calling | Glacier National Park (BC) |
| AB-2 | AB-2 | -117.39 | 53.48 | Calling | Hinton |
| AB-3 | AB-3 | -117.38 | 52.39 | Calling | Jasper National Park |
| BC-19 | BC-19 | -116.59 | 51.24 | Calling | Yoho National Park |
| AB-4 | AB-4 | -116.08 | 52.40 | Calling | Nordegg |
| AB-5 | AB-5 | -115.83 | 51.23 | Calling | Banff National Park |
| AB-6 | AB-6 | -115.56 | 52.50 | Calling | Clearwater County |
| AB-7a | AB-7 | -115.38 | 51.08 | Calling | Canmore |
| AB-7b | AB-7 | -115.38 | 51.08 | Calling | Canmore |
| AB-7c | AB-7 | -115.38 | 51.08 | Calling | Canmore |
| AB-7d | AB-7 | -115.38 | 51.08 | Calling | Canmore |
| AB-7e | AB-7 | -115.38 | 51.08 | Calling | Canmore |
| AB-8 | AB-8 | -115.26 | 51.97 | Calling | Clearwater County |
| AB-9 | AB-9 | -114.99 | 53.55 | Calling | Entwistle |
| BC-20 | BC-20 | -114.79 | 49.62 | Crown of the Continent | Corbin mines |
| BC-21 | BC-21 | -114.68 | 49.52 | Crown of the Continent | Corbin mines |
| AB-10 | AB-10 | -114.55 | 53.19 | Calling | Carnwood County |
| AB-11 | AB-11 | -114.05 | 49.05 | Crown of the Continent | Waterton Lakes National Park |
| AB-12 | AB-12 | -113.83 | 53.57 | Calling | Spruce Grove |
| AB-13 | AB-13 | -112.94 | 53.66 | Calling | Strathcona County |

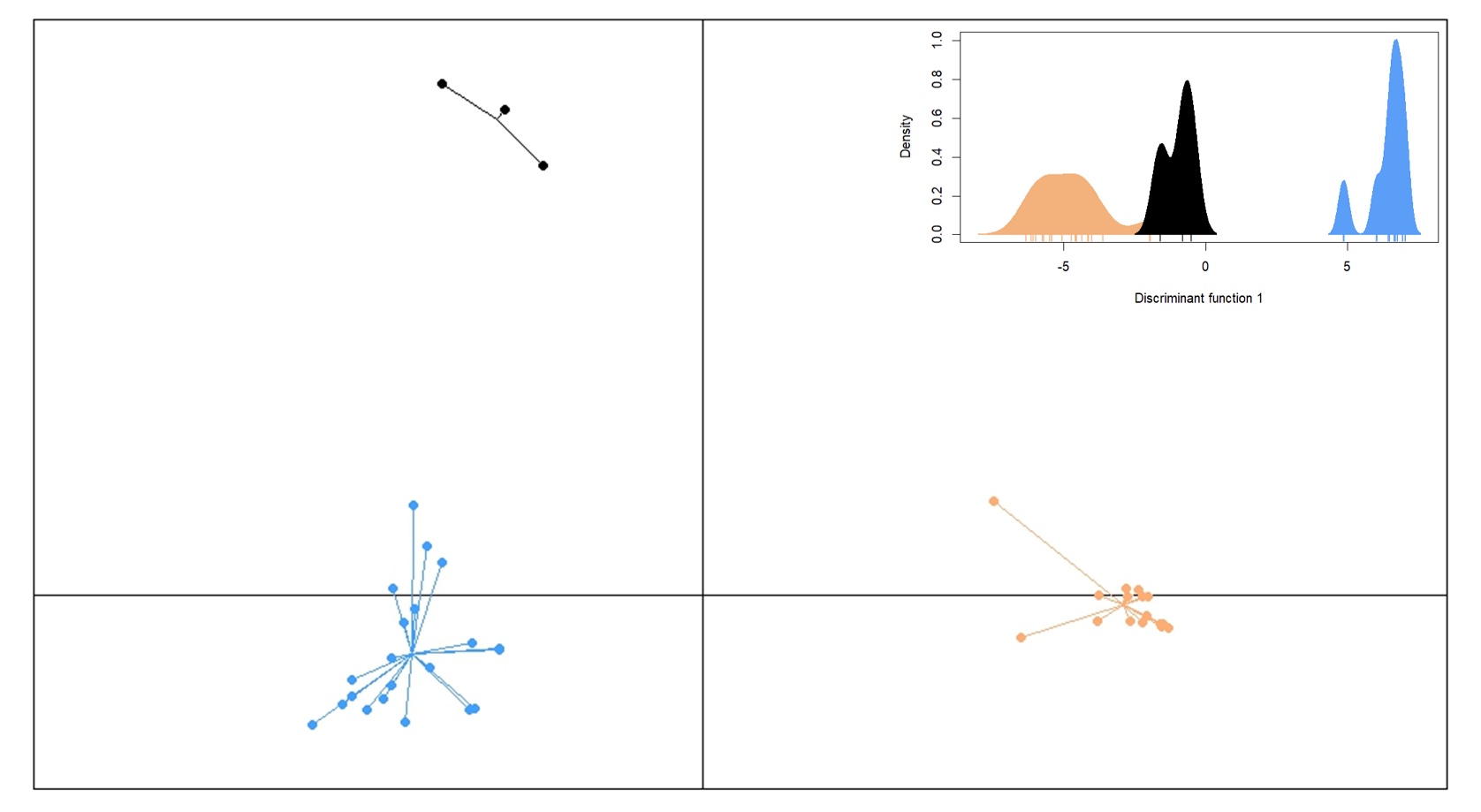

**Figure S1**. The Discriminant Analysis of Principal Components (DAPC) using 11,950 SNPs generated from 40 individuals across the Canadian range of western toads. Four principal components were retained for two discriminant functions. The top right inset shows the analysis contained within the first discriminant function as most of the information was within the first discriminant function. The colors are blue for the Non-Calling group, orange for the Calling group, and black for the Crown of the Continent group.

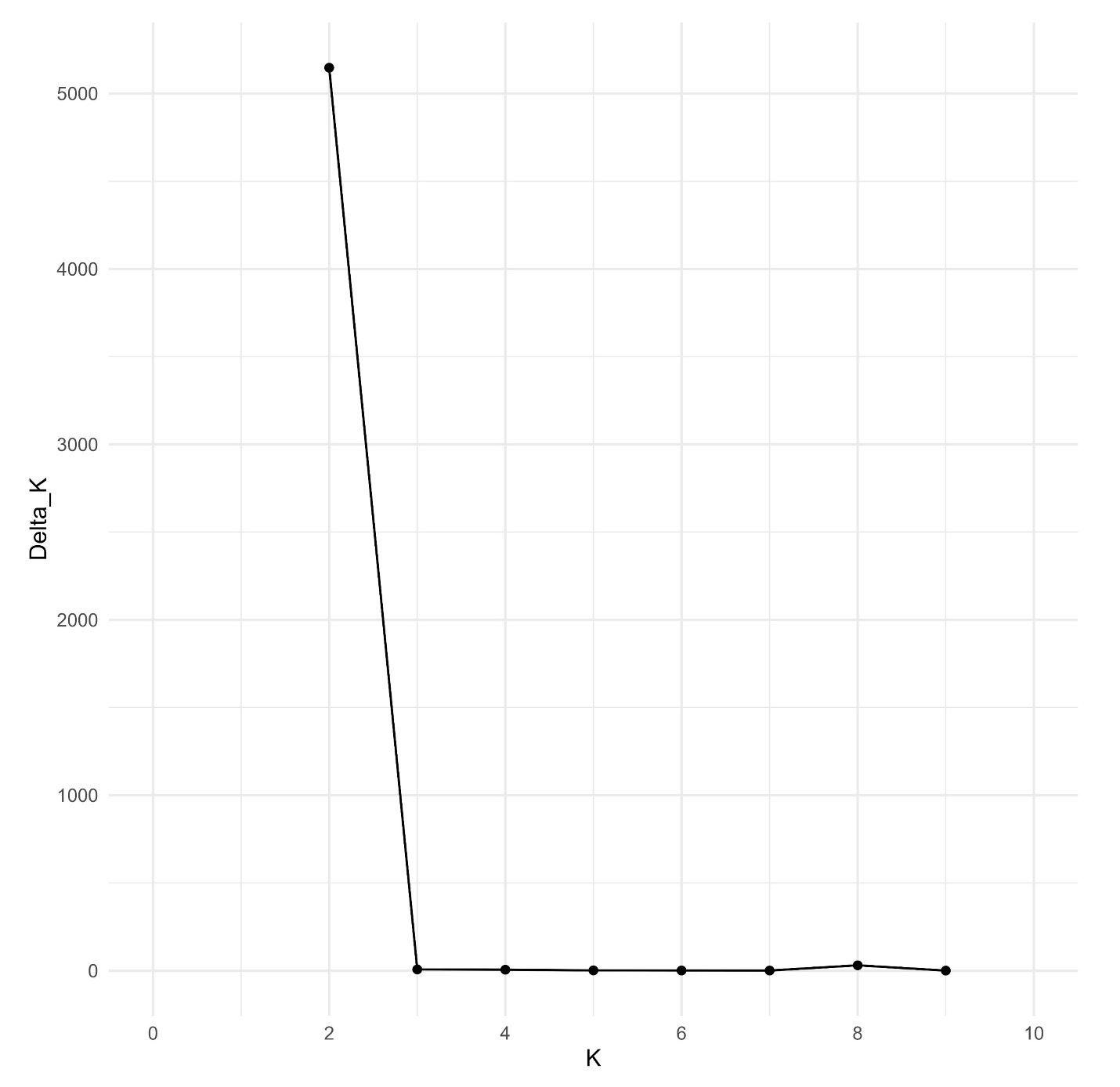

**Figure S2**. ΔK plot used to determine the optimal number of western toad populations (i.e. Evanno et al., 2005) in a STRUCTURE analysis of individuals from across the Canadian portion of the species’ range . STRUCTURE Harvester was used to generate this plot.

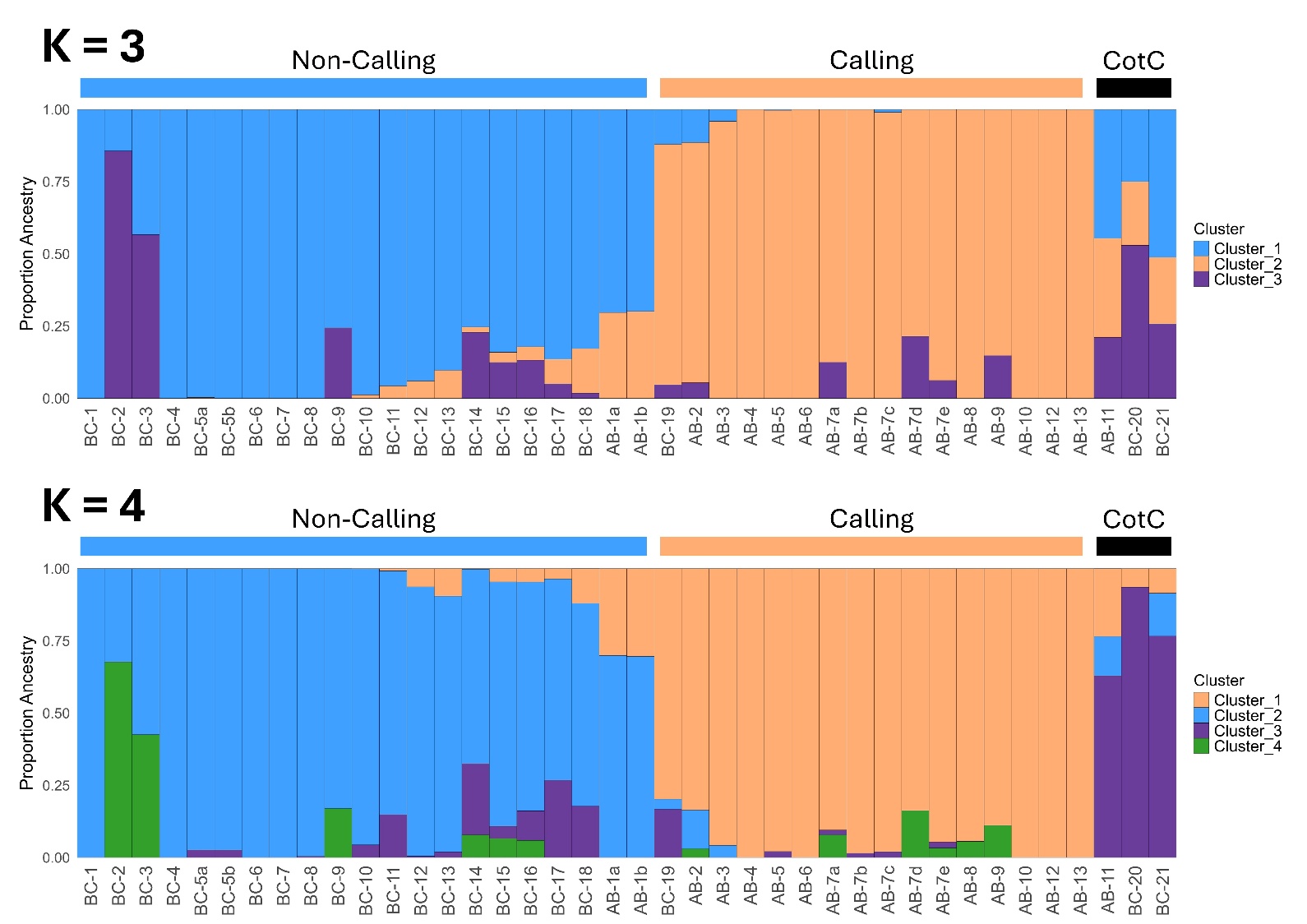

**Figure S3**. STRUCTURE results for (A) K=3 and (B) K=4 generated using 11,950 SNPs from 40 western toads sampled across the Canadian portion of the species’ range. The colored bars along the top of each plot represent the clustering of individuals found in the PCA in the main text (blue: non-calling, orange: calling, black: “Crown of the Continent” or CotC).

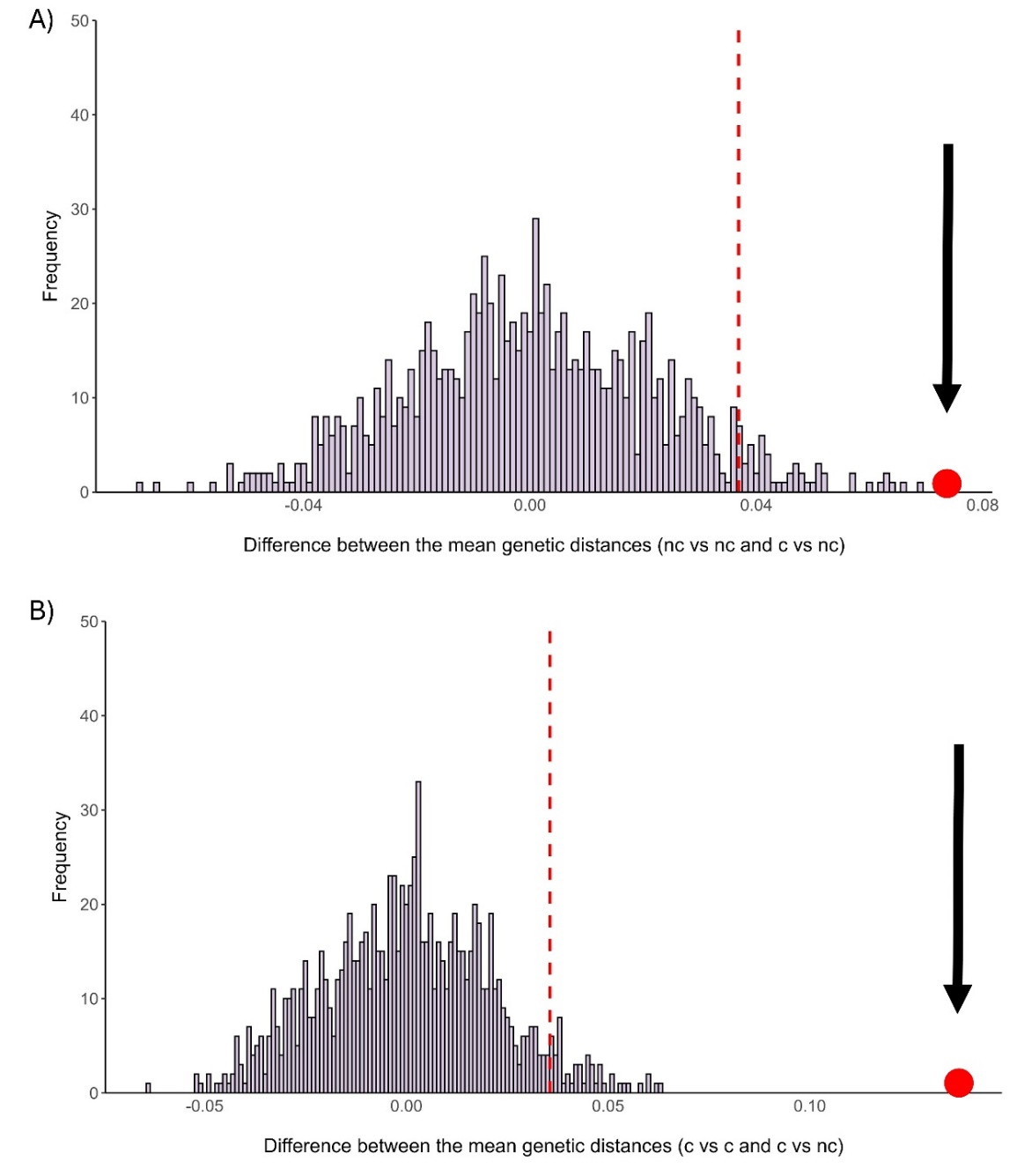

**Figure S4**. Testing the significance of differences in mean pairwise genetic distance (1 – proportion of shared alleles) of within-population versus between-population comparison types for western toads in Canada. (A) Pairwise differences among Calling individuals versus between Calling and Non-Calling individuals. (B) Pairwise differences among Non-Calling individuals versus between Calling and Non-Calling individuals. In both cases, individuals were within 150 km of each other. The histograms show the null distribution of mean pairwise differences generated by randomly assigning comparison type to each pair of individuals in the respective test. The dashed, red line in each plot show the 95^th^ quantile of differences, with the red point and black arrow indicating the observed mean pairwise. In both cases the observed value falls outside the 95^th^ quantile of the null distribution, suggesting observed differences between comparison types are greater than expected by chance.

**Table S2**. Details of model parameterization for ecological niche models generated for the Calling population and the Non-Calling population of western toads in Canada. Models were generated using Maxent and default parameter settings were used for other settings not specified.

| Population | Non-calling  model | Calling model |
| --- | --- | --- |
| No. input localities | 2852 | 215 |
| No. random background points | 5000 | 5000 |
| Feature classes* | LQPT | LQT |
| Regularization parameter | 0.5 | 1.5 |
| Maximum iterations | 5000 | 5000 |
| Training AUC | 0.73 | 0.73 |
| 10% omission threshold value to consider a cell as suitable | 0.379 | 0.365 |

*L = linear, Q = quadratic, P = product, T = threshold

**Table S3**. Percent contribution and permutation importance of the eight climatic variables included in the Maxent models for the Calling population and the Non-calling population of western toads in Canada.

| **Variable** | **Importance** | **Non-calling model** | **Calling model** |
| --- | --- | --- | --- |
| Mean temperature of the coldest month (MCMT) | Percent contribution | 9.7 | 18.7 |
|  | Permutation importance | 8 | 8.1 |
| Summer (June to August) precipitation (mm) (PPT_sm) | Percent contribution | 10.4 | 28.9 |
|  | Permutation importance | 4.8 | 9.7 |
| Julian date on which the frost-free period begins (bFFP) | Percent contribution | 2 | 38 |
|  | Permutation importance | 2.8 | 39.3 |
| Summer heat moisture index (SHM) | Percent contribution | 6.7 | 2.8 |
|  | Permutation importance | 14.4 | 5.7 |
| Precipitation as snow (mm) (PAS) | Percent contribution | 44 | 4.4 |
|  | Permutation importance | 5.6 | 5.4 |
| Spring (March to May) precipitation (mm) (PPT_sp) | Percent contribution | 6.8 | 5.4 |
|  | Permutation importance | 27.5 | 7 |
| Degree-days below 18 degrees Celsius (DD_18) | Percent contribution | 15.7 | 0.6 |
|  | Permutation importance | 33.7 | 18.4 |
| Mean annual relative humidity (%) (RH) | Percent contribution | 4.8 | 1.2 |
|  | Permutation importance | 3.2 | 6.5 |

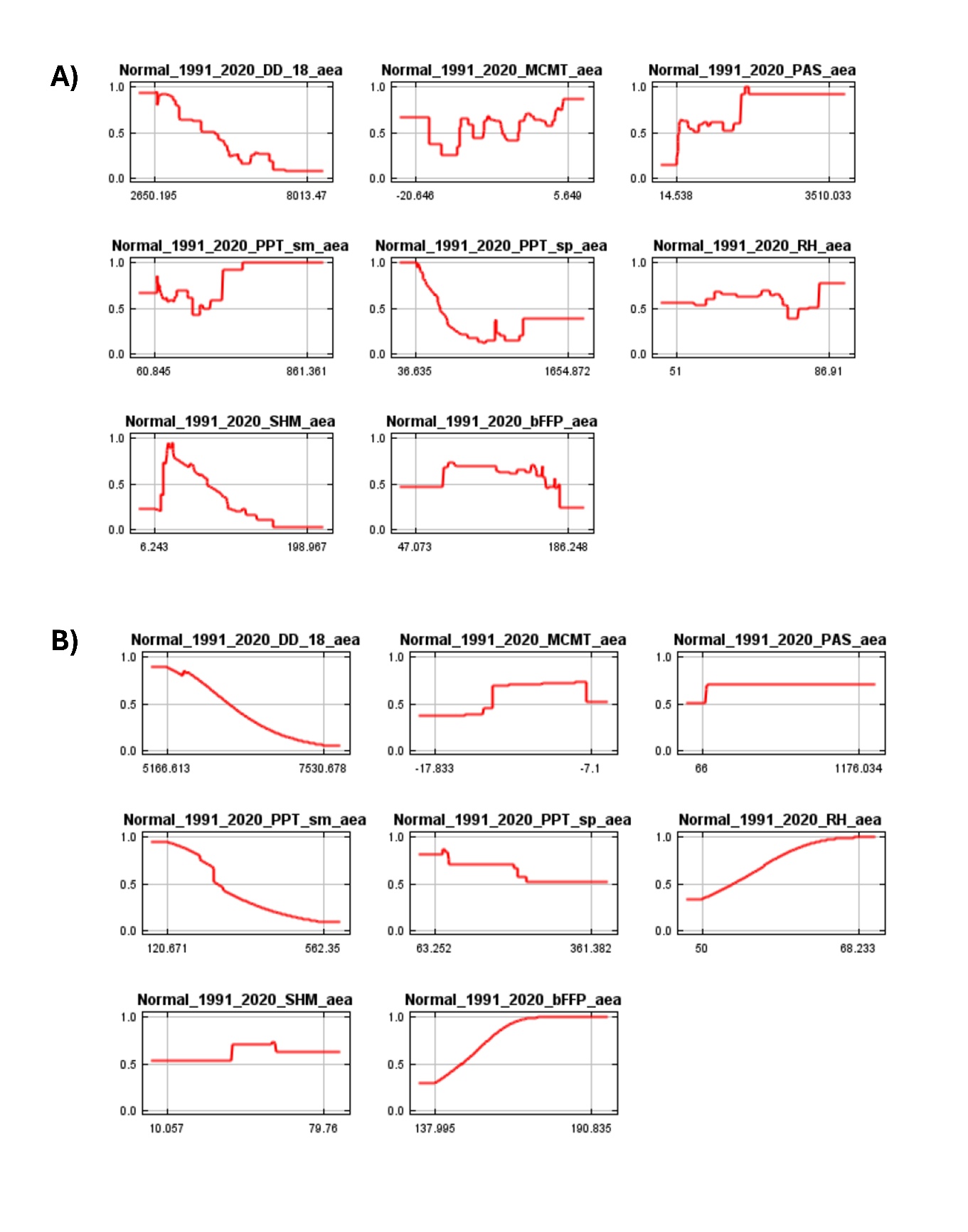

**Figure S5**. Response curves for each climatic variable included in the Maxent models for the A) Non-Calling population and the B) Calling population.

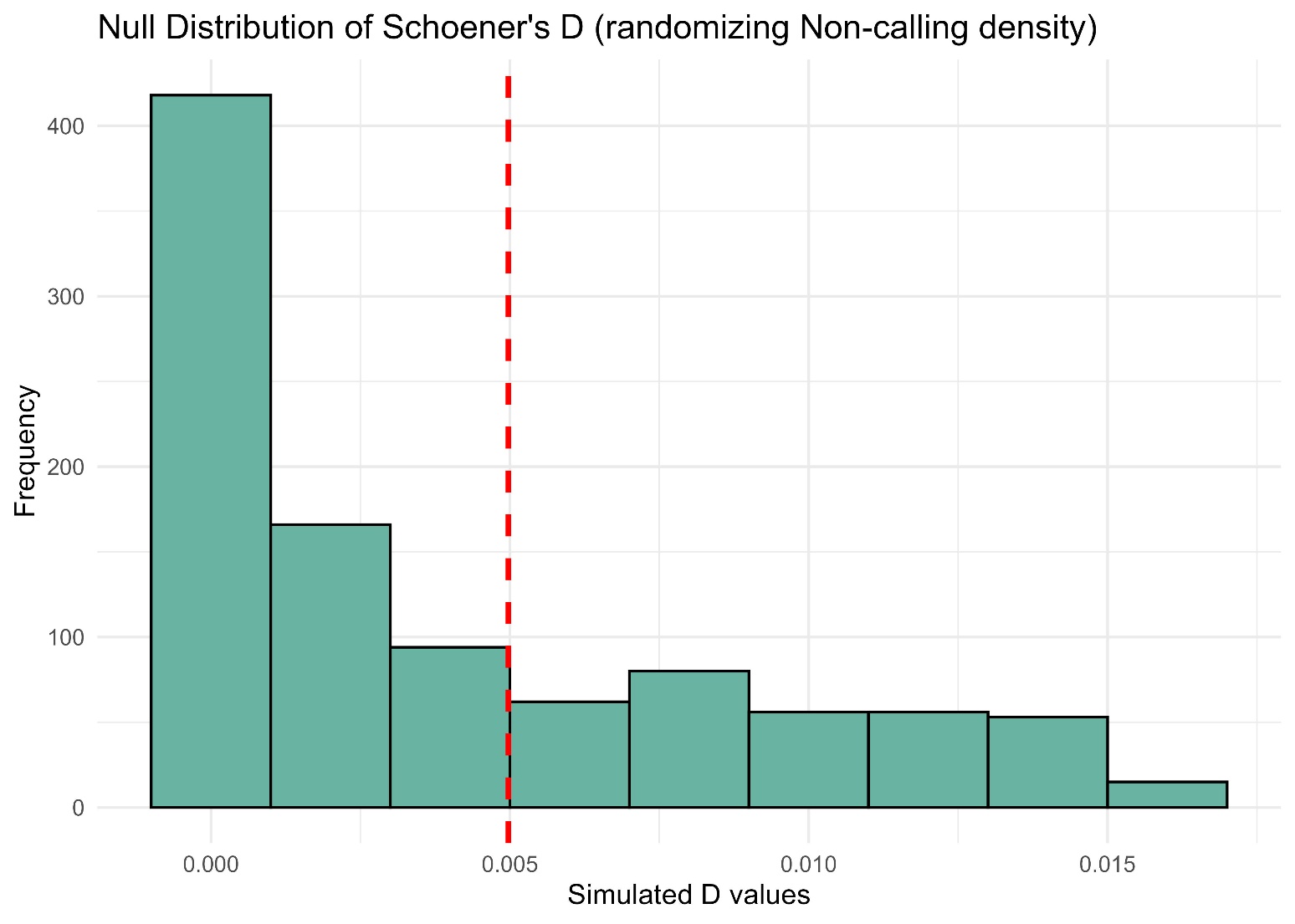

**Figure S6**. Simulated Schoener’s D values via 1000 randomizations to assess the statistical significance of niche similarity using the ecospat.niche.similarity.test() in the R package ecospat (Broennimann et al., 2025). The red dashed line is the observed niche similarity (Schoener’s D = 0.005, p-value = 0.309) when the Non-Calling population’s niche is randomized.

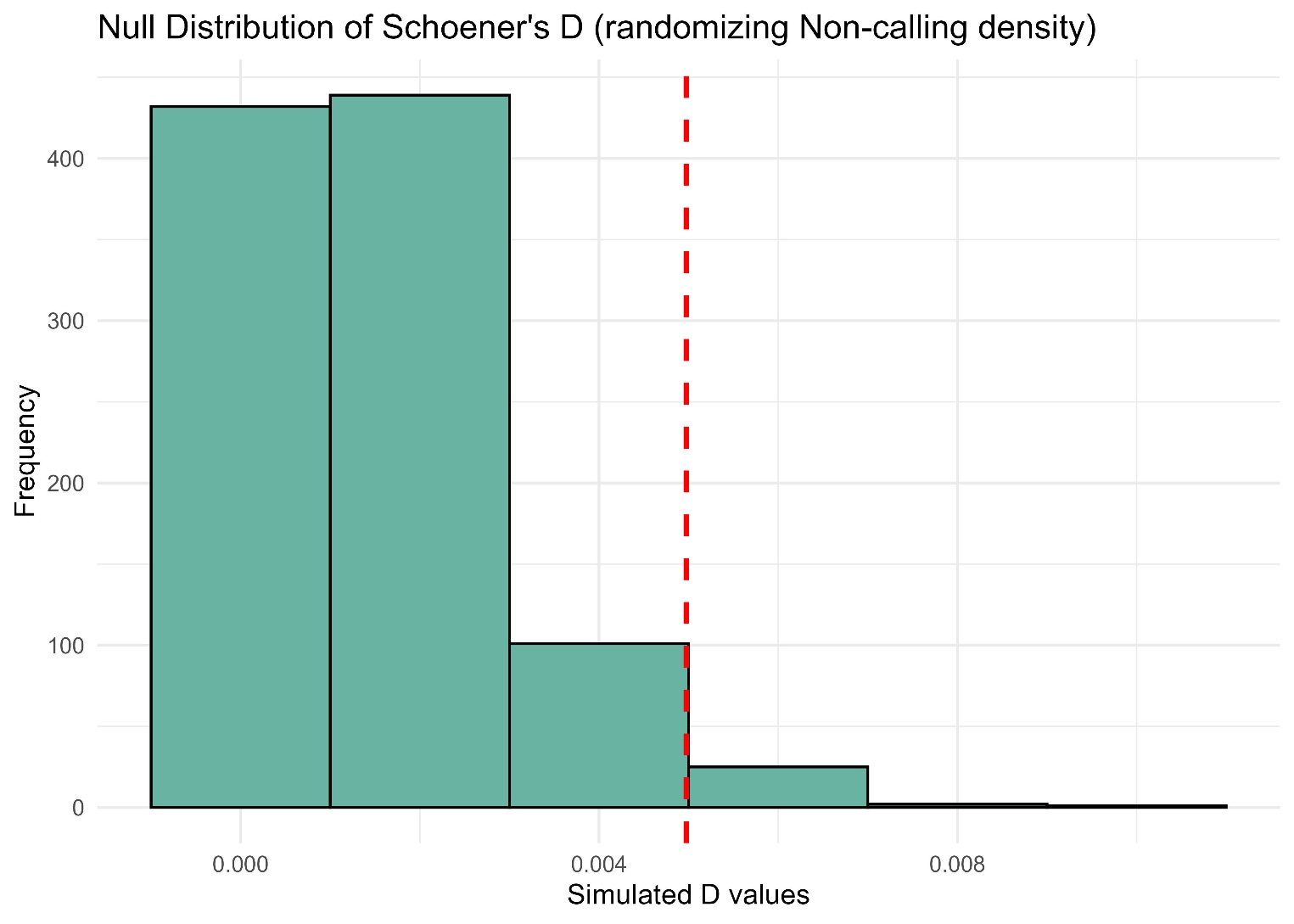

**Figure S7**. Simulated Schoener’s D values via 1000 randomizations to assess the statistical significance of niche similarity using the ecospat.niche.similarity.test() in the R package ecospat (Broennimann et al., 2025). The red dashed line is the observed niche similarity (Schoener’s D = 0.005, p-value = 0.048) when the Calling population’s niche is randomized.
